# Conservation of *cis*-regulatory codes over half a billion years of evolution

**DOI:** 10.1101/2024.11.13.623372

**Authors:** Yohey Ogawa, Yu Liu, Connie A. Myers, Ala Morshedian, Gordon L. Fain, Alapakkam P. Sampath, Joseph C. Corbo

## Abstract

The identification of homologous cell types across species represents a crucial step in understanding cell type evolution. The retina is particularly amenable to comparative analysis because the basic morphology, connectivity, and function of its six major cell classes have remained largely invariant since the earliest stages of vertebrate evolution. Here, we show that the retina’s highly conserved cellular architecture is mirrored by deep conservation of the underlying *cis*-regulatory codes that control gene expression. We use comparative single-cell chromatin accessibility analysis of lamprey, fish, bird, and mammalian retinas— representing over half a billion years of evolutionary divergence—to demonstrate cross-species conservation of *cis*-regulatory codes in all six retinal cell classes. This conservation persists despite extensive turnover of *cis*-regulatory regions between distant species. Conservation manifests as the clustering of multiple distinct high-affinity transcription factor (TF) binding sites toward the center of cell-class-specific open chromatin regions with little cross-species preservation of higher-order syntax. Hierarchical clustering of machine-learning models of retinal *cis*-regulatory codes from diverse species recovers six clusters corresponding to the six retinal cell classes. Thus, the retina’s cellular *Bauplan* is controlled by *cis*-regulatory codes which predate the divergence of extant vertebrates and persist despite nearly complete enhancer turnover.

## Introduction

The evolutionary origin of cell types is a subject of enduring fascination^1–4^. To infer the existence of specific cell types in the common ancestor of extant vertebrates—a species that lived ∼560 million years ago—it is necessary to compare homologous cell types between the two most evolutionarily distant vertebrate taxa: the jawless fishes (i.e., lampreys and hagfishes) and the jawed vertebrates (cartilaginous and bony fishes, amphibians, reptiles, birds, and mammals). The vertebrate retina is an ideal system for inferring cellular and molecular features that existed in the common vertebrate ancestor, because its basic features—cell classes, connectivity patterns, and function—are remarkably conserved among all vertebrate taxa^5–7^, including between jawed and jawless species^8–10^ (Fig. 1a, b). Nearly all vertebrate retinas contain six major cell classes (photoreceptors, bipolar cells, horizontal cells, amacrine cells, ganglion cells, and Müller glia)^7^. Each cell class— with the exception of Müller glia— consists of multiple closely related ‘sister’ cell types—expressing divergent sets of effector genes but retaining many shared transcriptional regulators—which arose via duplication and divergence from a single ancestral cell type^1,2,11^. In the course of evolution, individual vertebrate species have expanded or contracted the number of cell types within each cell class to adapt to specific light environments or lifestyles^6,12,13^. Thus, vertebrate retinas display remarkable cell type diversity couched within an evolutionarily stable framework of six invariant cell classes.

**Fig. 1.**
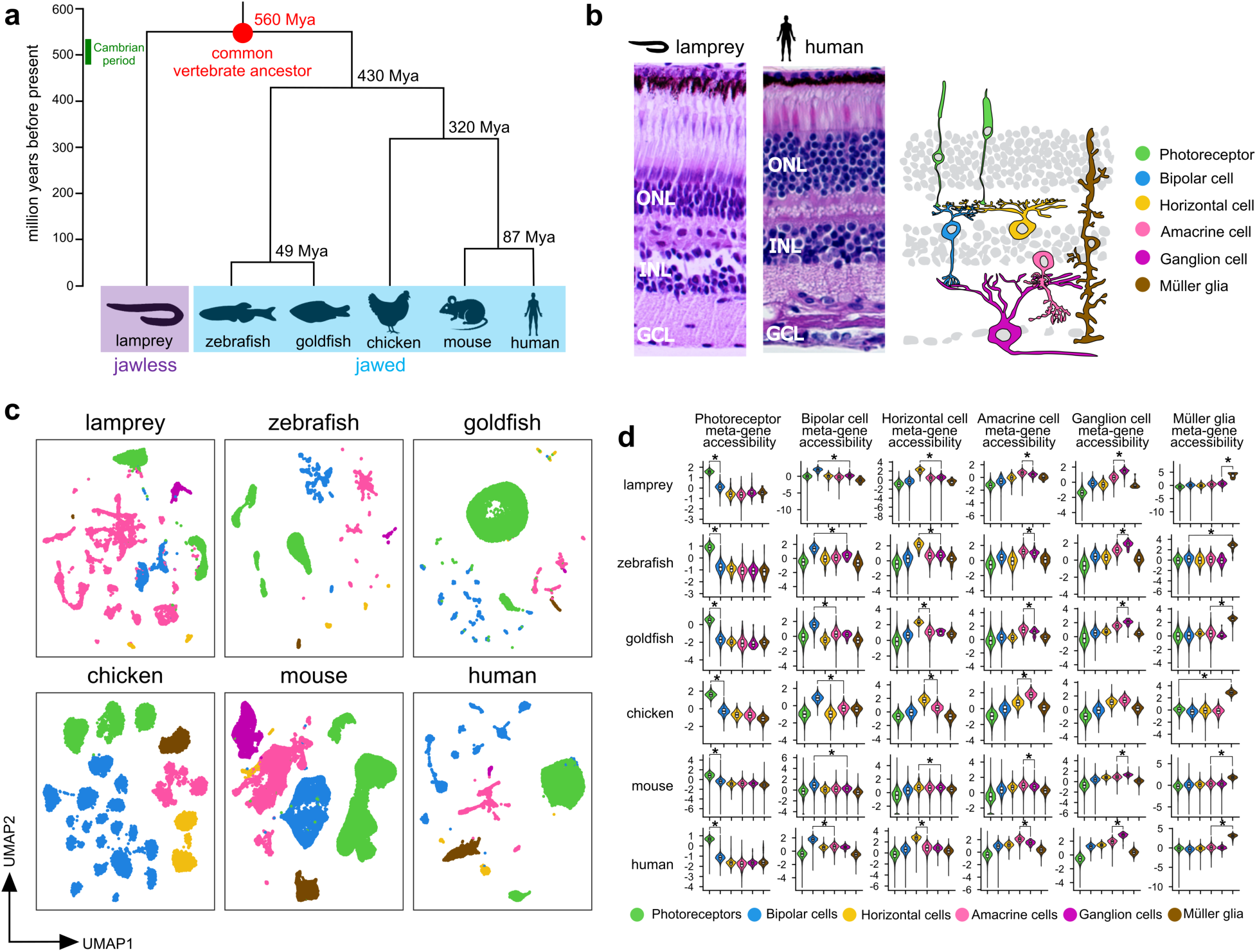
Single-nucleus chromatin accessibility profiles for six retinal cell classes in six vertebrate species. **a,** A phylogenetic tree of the vertebrate species utilized in this study. Divergence times are estimated based on a published database^62^. **b,** (left) H&E-stained sections of lamprey and human retina. (right) Schematic of the six major retinal cell classes in vertebrates. ONL, outer nuclear layer; INL, inner nuclear layer; GCL, ganglion cell layer. **c,** Single-cell ATAC-seq profiles of the six indicated species. Retinal cell classes were identified by clustering analysis and visualized in two-dimensional space using the Uniform Manifold Approximation and Projection (UMAP) method. Cell classes are color-coded as in **b**. **d,** Violin plots showing the average chromatin accessibility of cell class-enriched meta-genes generated from sets of evolutionarily conserved marker genes for each cell class in each species (see Methods). The rows are grouped by species for each snATAC-seq data set, and the columns are grouped by cell class. Cell-class meta-gene enrichment was determined by comparing the cell class exhibiting the highest score for the meta-gene with the second-highest scoring cell class. An asterisk indicates an adjusted p-value <0.05 (Wilcoxon rank sum test followed by Bonferroni correction).

Cell class-and type-specific transcriptomes are determined by the action of transcriptional regulatory networks, which consist of hierarchical cascades of TFs that bind to cognate binding sites within *cis*-regulatory elements (i.e., enhancers and promoters) to regulate gene expression and determine cell type identity^14,15^. A ‘*cis*-regulatory code’ or ‘grammar’ is the particular combination and arrangement of TF binding sites within *cis*-regulatory elements that drives expression in a specific cell type or class. In the present study, we sought to determine whether the *cis*-regulatory codes governing retinal class-specific gene expression are conserved across vertebrates, including between jawed and jawless species. Indeed, we find that the architectural invariance of the vertebrate retina is mirrored by deep conservation of the underlying *cis*-regulatory codes and that these codes emerged in the common ancestor of extant vertebrates more than half a billion years ago.

## Results

### Single-cell chromatin accessibility profiling of vertebrate retinas

To determine the evolutionary antiquity of the *cis*-regulatory codes that govern gene expression in the vertebrate retina, we carried out a systematic analysis of TF binding sites in the retinal cell class-enriched open chromatin regions of six diverse vertebrate species (Fig. 1). These species inhabit a wide range of photic environments and have correspondingly evolved divergent retinal cell type inventories^7,16^. Thus, it is impossible to define one-to-one homology relationships for individual cell types across all species. We therefore focused our analysis on cell class-specific *cis*-regulatory codes. To accomplish this task, we acquired published retinal single-cell gene expression profiling (scRNA-seq) and single-nucleus chromatin accessibility (snATAC-seq) data from two teleost fishes (zebrafish and goldfish) and two placental mammals (mouse and human). To broaden our phylogenetic sampling, we additionally conducted single-cell analyses on retinas from chicken (*Gallus gallus*) and sea lamprey (*Petromyzon marinus*), a jawless species. We examined these six datasets—generated by distinct protocols and from diverse sources—using either *Signac/Seurat* or *ArchR*, depending on whether the data were generated by multiome (snRNA-seq + snATAC-seq) or snATAC-seq analysis, respectively. After initial pre-processing to remove low-quality cells, we performed dimensionality reduction followed by shared nearest neighbor modularity optimization-based clustering. For each species, we assigned clusters to one of the six retinal cell classes, either based on the expression of known class-specific marker genes in those species for which multiome data was available (lamprey, zebrafish, and human), or based on chromatin openness at the promoters of class-specific marker genes in those species for which multiome data was not available (goldfish, chicken, and mouse). We removed from the analysis any residual clusters which could not be assigned to one of the major retinal cell classes. In this way, we identified clusters corresponding to each of the six retinal cell classes in all six species, with the exception of chicken ganglion cells, which were absent from the snATAC-seq dataset, though present in scRNA-seq data (Fig. 1c; Supplementary Fig. 1).

Next, we sought to determine whether our cluster annotations are reflective of shared patterns of chromatin openness at class-enriched gene loci. To achieve this goal, we devised a quantitative measure of class-specific chromatin openness for each of the six retinal cell classes. First, we used scRNA-seq data to define evolutionarily conserved class-specific ‘meta-genes’ consisting of a set of genes that were differentially expressed across cell classes and had a high class-specificity index in lamprey and three or more of the jawed species (see Methods). We then measured chromatin openness over the promoter and gene body of each gene in the meta-gene and aggregated the values to create a single ‘meta-gene openness’ score for each of the six cell classes in each of the six species (except for chicken ganglion cells; see above). We found that in all comparisons the meta-gene openness score was highest for the expected cell class (Fig. 1d). These findings validate our cluster annotations and confirm that class-specific signatures of chromatin openness are shared across all six species.

### Extensive enhancer turnover during vertebrate evolution

Next, we wished to determine if retinal class-specific *cis*-regulatory elements show sequence-level conservation across species. *Cis*-regulatory elements typically occur in chromatin regions that are selectively open in the cell type(s) in which the element is active. For example, we previously showed extensive overlap between photoreceptor-enriched open chromatin regions and the location of photoreceptor-specific enhancers and promoters across the mouse genome^17,18^. We therefore decided to use class-enriched open chromatin regions (OCRs) as a surrogate for class-enriched *cis*-regulatory elements in the present analysis. To quantify the extent of sequence-level conservation of class-enriched OCRs across species, we used single-cell data to create pseudo-bulk ATAC-seq profiles for each of the six retinal cell classes in five species: lamprey, zebrafish, chicken, mouse, and human (except for chicken ganglion cells; see above). We identified class-enriched OCRs for each retinal cell class in each species using a test of differential chromatin accessibility (Supplementary Fig. 2; see Methods). We then employed the UCSC Genome Browser’s *LiftOver* utility to map the union of each species’ class-enriched OCRs onto all other vertebrate reference genomes for which pre-computed reciprocal best-hit whole-genome alignment files were publicly available. In this way, we quantified ‘alignability’ as the percentage of a species’ class-enriched OCRs which could be aligned with the genome of the target species (see Methods for details). We found that sequence alignability of retinal class-enriched OCRs progressively decayed with evolutionary distance such that beyond ∼400 million years (Mys) fewer than 3% of OCRs were alignable with the target genome (Fig. 2; Supplementary Table 1). For example, the average alignability at 430 Mys (the distance between ray-finned fishes and amniotes) was 1.35%, while the average alignability at 563 Mys (the distance between jawed and jawless species) was 0.52%. We therefore infer that vertebrate *cis*-regulatory element turnover is extensive at great evolutionary distances.

**Fig. 2.**
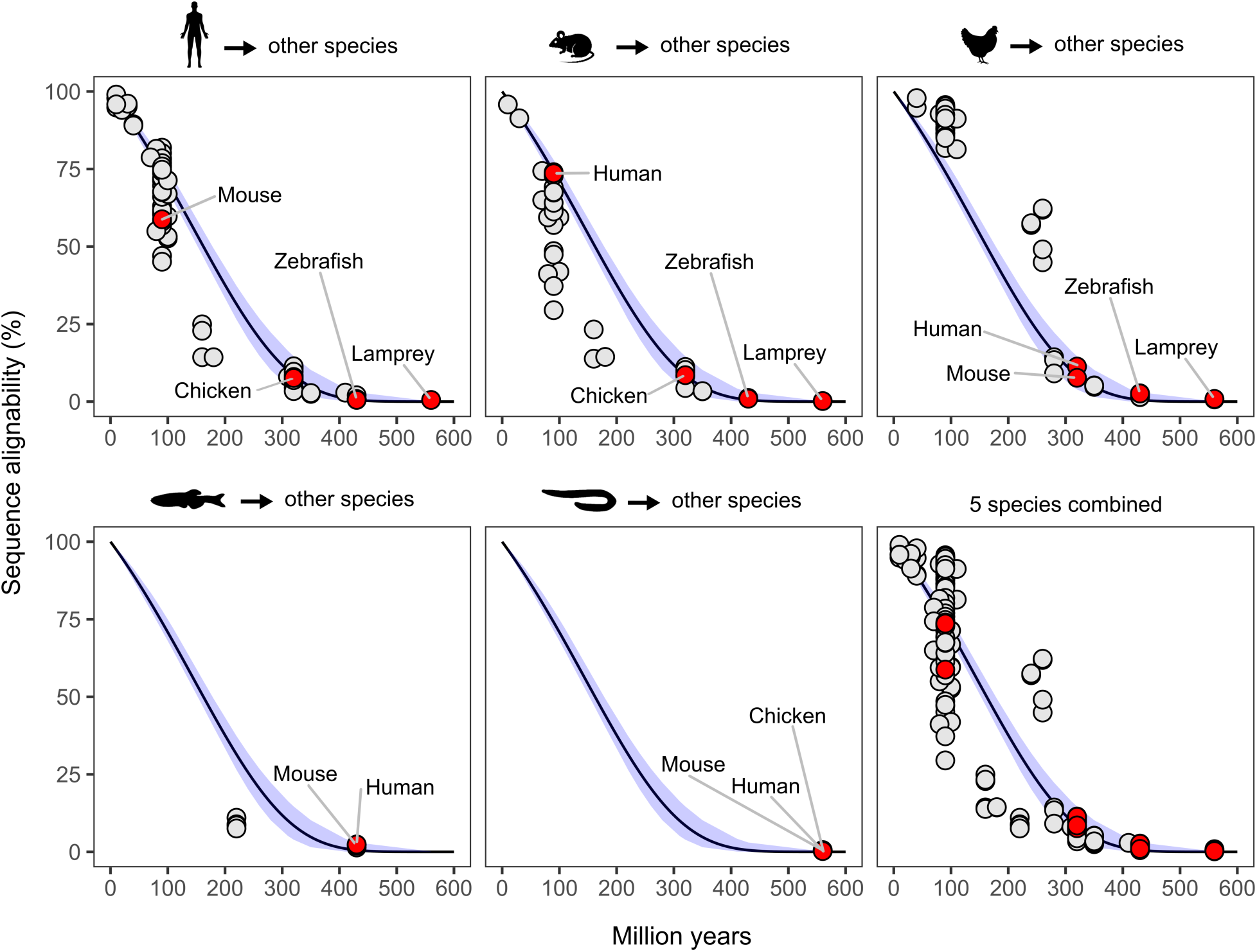
Nearly complete sequence turnover of retinal cell class-enriched open chromatin regions after 400 million years of evolution. Retinal cell class-enriched chromatin regions in five query species (human, mouse, chicken, zebrafish, and lamprey) were mapped onto the genomes of diverse vertebrate species (also see Supplementary Table 1). The X-axis represents the evolutionary divergence times between query and target species according to a published database^62^. The Y-axis indicates the percent of query sequences that can be aligned to the target genome. The decay of sequence alignability over evolutionary time was modeled with the Gompertz equation using divergence time as a variable. The 95% confidence intervals measured by bootstrap resampling (see Methods) are shown as blue shading. The query species, when used as targets, are labeled and highlighted in red.

To model the decline in OCR alignability over time, we fitted the data with a Gompertz equation, which is often used to model the growth or decay of populations. We observed close agreement with the model at short (i.e., <50 Mys) and long (>280 Mys) evolutionary distances but found major deviations from the model at middle distances (50-280 Mys) (Fig. 2; Supplementary Table 1). The greatest downward deviations—indicative of more extensive turnover than predicted by the model—were observed in mouse/human-to-mammal comparisons, particularly at ∼90 Mys (i.e., mouse/human-to-placental comparisons), 160 Mys (mouse/human-to-marsupial), and 180 Mys (mouse/human-to-monotreme) (Fig. 2). These deviations are likely attributable to accelerated rates of evolution in certain mammalian clades^19,20^. Conversely, we observed unexpectedly low rates of OCR turnover in chicken-to-bird comparisons (∼90 Mys) and chicken-to-alligator/turtle comparisons (240-260 Mys), while chicken-to-lizard/snake comparisons (280 Mys) largely agreed with the model (Fig. 2). Despite wide variation in the rates of alignability at middle evolutionary distances, both the data and the model suggest nearly complete (i.e., ∼99.5%) evolutionary turnover of retinal class-specific *cis*-regulatory elements beyond 500 million years.

### Deep conservation of retinal *cis-*regulatory codes

We and others observed conserved patterns of expression of class-specific genes despite extensive turnover of the *cis*-regulatory elements controlling their expression (Supplementary Table 2)^7,9^. We therefore hypothesized that the underlying *cis*-regulatory codes might be conserved despite an absence of linear sequence conservation. To test this idea, we undertook a detailed comparative analysis of retinal class-specific *cis*-regulatory codes. The fundamental building blocks of a *cis*-regulatory code are TF binding sites. Thus, as a first step toward elucidating the retinal codes, we used *HOMER*^21^, a TF binding site motif discovery algorithm, to comprehensively identify motifs enriched within 201-bp regions centered on the summits of class-specific OCRs of six species. All existing motif databases derive from in-depth study of a small number of species. Thus, to avoid biases that might be introduced by focusing on ‘known’ motifs, we used *HOMER* to detect *de novo* motifs. *HOMER* identifies enriched motifs by comparing an ‘experimental’ set of target sequences with a set of control sequences. We therefore used class-enriched OCRs as our experimental dataset and a set of OCRs broadly open across multiple cell classes as our controls (Fig. 3a). *HOMER* generates a list of motifs in the form of a position probability matrix accompanied by an optimal detection threshold to maximize the enrichment of the motif in the target sequences. To ensure detection of relatively low-frequency but functionally important motifs, we retained all motifs whose statistical significance of enrichment was <10^-10^ (calculated using the binomial distribution) and which were present in >2% of the cell-class enriched OCRs. We analyzed a total of 35 datasets (i.e., six species × six cell classes, except for chicken ganglion cells), identifying a median of 47 *de novo* motifs in each dataset, with a minimum of eight motifs observed for lamprey Müller glial cells, likely due to the small number of these cells in our dataset (Supplementary Table 3).

**Fig. 3.**
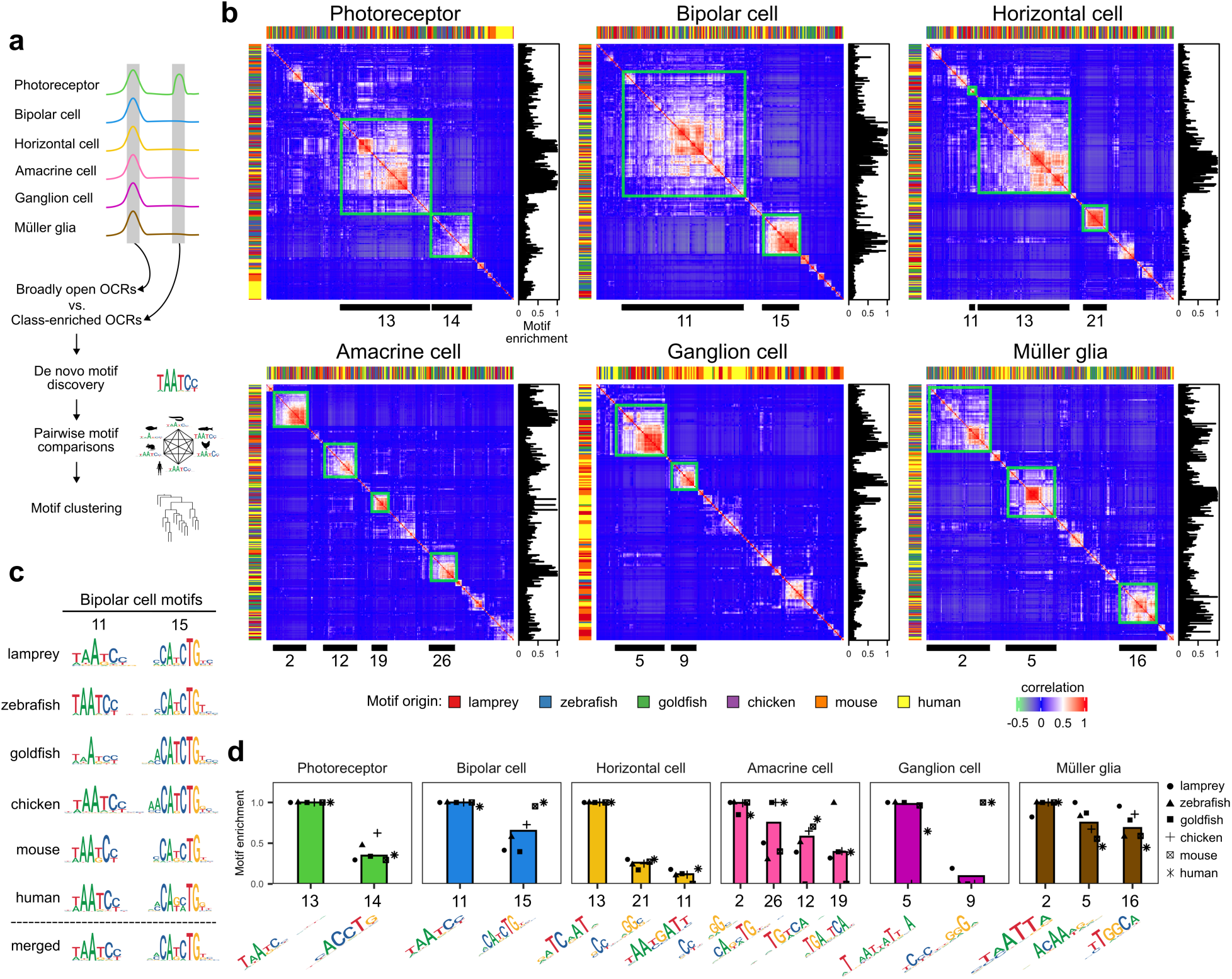
Multiple *cis*-regulatory motifs for each retinal cell class are conserved between jawed and jawless species. **a,** Schematic showing the methodology used for the identification of cell class-enriched open chromatin regions, the discovery of sequence motifs, and motif clustering. **b,** Heatmaps showing motif-similarity correlation values for all pairs of significantly enriched *de novo* motifs from six species. The motif-pair values were hierarchically clustered to reveal families of related motifs (see Methods). Evolutionarily conserved motif families—as defined in the main text—are enclosed by green boxes and indicated by numbered black bars at the bottom of each heatmap. The species of origin for each motif is indicated by color-coding across the top and left sides of each heatmap. ‘Motif enrichment’ is the normalized statistical significance of motif enrichment in each species (i.e., the most enriched motif in each species has a motif enrichment = 1). **c,** Representative examples of two evolutionarily conserved motifs (EC motifs) in bipolar cells. The sequence logo for the most significantly enriched motif in each species is shown, along with the logos for a ‘merged’ motif representing the average of the six species motifs. For the full set of EC motifs, see **Supplementary Fig. 3. d**, The median of the normalized statistical significance of motif enrichment for each of the 16 *de novo* motifs is shown (see Methods). The normalized motif enrichment value for each of the individual species is also presented. The sequence logos at the bottom represent the merged motifs.

Next, we sought to compare class-specific motifs across species to determine if diverse vertebrates utilize a shared set of motifs in each retinal cell class. To accomplish this goal, we used *Tomtom*^22^ to conduct pairwise motif comparisons and then hierarchically clustered motifs based on their similarity scores. *HOMER* often identifies multiple related motifs; thus the several dozen *de novo* motifs discovered for a given cell class in an individual species may correspond to a smaller set of truly distinct motifs. By including all identified motifs for a given cell class in our hierarchical clustering, we can both delineate intraspecific motif redundancy and identify interspecific similarities in motif inventory.

Visual inspection of motif similarity matrices for each of the six retinal cell classes reveals multiple well-defined clusters of motifs for each cell class (Fig. 3b). Motifs in these clusters typically exhibit some of the highest ‘motif enrichment’ scores (Fig. 3b; see Methods). Additionally, some motif clusters are quite large, encompassing as much as ∼35-50% of the motifs in a given cell class (e.g., in photoreceptors, bipolar cells, and horizontal cells), underscoring the presence of motif redundancy in the *HOMER* outputs. Importantly, within individual clusters we typically find motifs from multiple species, indicative of cross-species conservation of motifs. To define discrete motif clusters likely corresponding to individual conserved motifs, we truncated the motif dendrogram (obtained by hierarchical clustering) at a height of 0.9 (equal to one minus the Pearson correlation coefficient of motif similarity values). We then designated a motif cluster as ‘evolutionarily conserved’ if it included motifs from both jawless (i.e., lamprey) and three or more jawed species, and if the median of the Pearson correlation coefficient among the motif similarities of the most enriched motifs from each species within a cluster was greater than 0.5 (Fig. 3b and 3c; see Methods). For each evolutionarily conserved motif cluster, the motifs with the highest ‘motif enrichment’ score from each species were aggregated into a single ‘merged motif’ (see Methods), which we designated as an ‘evolutionarily conserved motif’ (EC motif) (Fig. 3c and Supplementary Fig. 3). In this way, we identified a total of 16 EC motifs, with two to four motifs in each retinal cell class (Fig. 3b, d). We propose that these motifs formed part of the retinal *cis*-regulatory codes of the most recent common ancestor of extant vertebrates based on their enrichment in both jawed and jawless species.

The close similarity of the species-specific position probability matrices used to create merged EC motifs suggests that these motifs are bound by homologous TFs with very similar DNA-binding preferences across species. To nominate TFs likely to bind these EC motifs, we used *Tomtom* to compare EC motifs to known motifs in *HOCOMOCO*^23^, a curated database of mouse and human TF binding site motifs. We found that all EC motifs showed highly significant matches to one or more motifs in the database (Supplementary Fig. 3 and Supplementary Table 4). For example, the most enriched EC motifs in photoreceptors (PH_13) and bipolar cells (BC_11) are very similar to each other and closely match paired-type ‘K50’ homeodomain binding sites (‘K50’ denoting the presence of lysine at position 50 of the homeodomain) bound by CRX and/or OTX2 in the *HOCOMOCO* database. These TFs are both expressed in mammalian photoreceptor and bipolar cells and play critical roles in controlling development and gene expression in these cell classes^24,25^. Indeed, most of the EC motifs show matches to binding sites of mammalian TFs previously shown to play key roles in regulating gene expression in their respective cell classes (Supplementary Table 4)^26,27^.

We postulated that the non-mammalian species in our study likely also express TFs in their respective cell classes with similar binding preferences to those of their mammalian counterparts. To test this idea, we mapped the candidate mammalian TFs onto their closest homologs in the non-mammalian species using *OrthoFinder*^28^. We then intersected the resultant TF orthology groups with lists of differentially expressed genes obtained from single-cell RNA-seq profiling of retinas from each of the six species. For 12 of the 16 EC motifs, we were able to identify cognate orthologous TFs whose expression was enriched in the corresponding cell class in jawless and four or more jawed species (Supplementary Table 4; see Methods). This finding suggests that cross-species conservation of class-enriched motifs is paralleled by cross-species conservation of cognate TF expression.

Next, we sought to systematically determine which features of class-specific *cis*-regulatory grammar are conserved across species. *Cis*-regulatory grammar can be subdivided into two components: ‘vocabulary’ consisting of the occurrence, affinity, and location of individual motifs within an OCR; and ‘syntax’ comprising the co-occurrence, spacing, and relative orientation of pairs of motifs in an individual OCR (Fig. 4a). So far we have identified significant class-specific enrichment of 16 EC motifs (Fig. 3). We next determined the spatial distribution of these motifs within class-enriched OCRs and quantified their position-weight-matrix (PWM) scores, a surrogate measure of binding affinity. We found that most motifs display a Gaussian-like distribution of enrichment conserved across species with a peak centered on the OCR summit (Fig. 4b). PWM scores also peaked at the OCR summit (Fig. 4b). In most cases, both motif enrichment and PWM scores declined monotonically with distance from the summit, approaching baseline levels around ±100 bp. These results indicate that features of *cis*-regulatory vocabulary are largely shared across motifs and species.

**Fig. 4.**
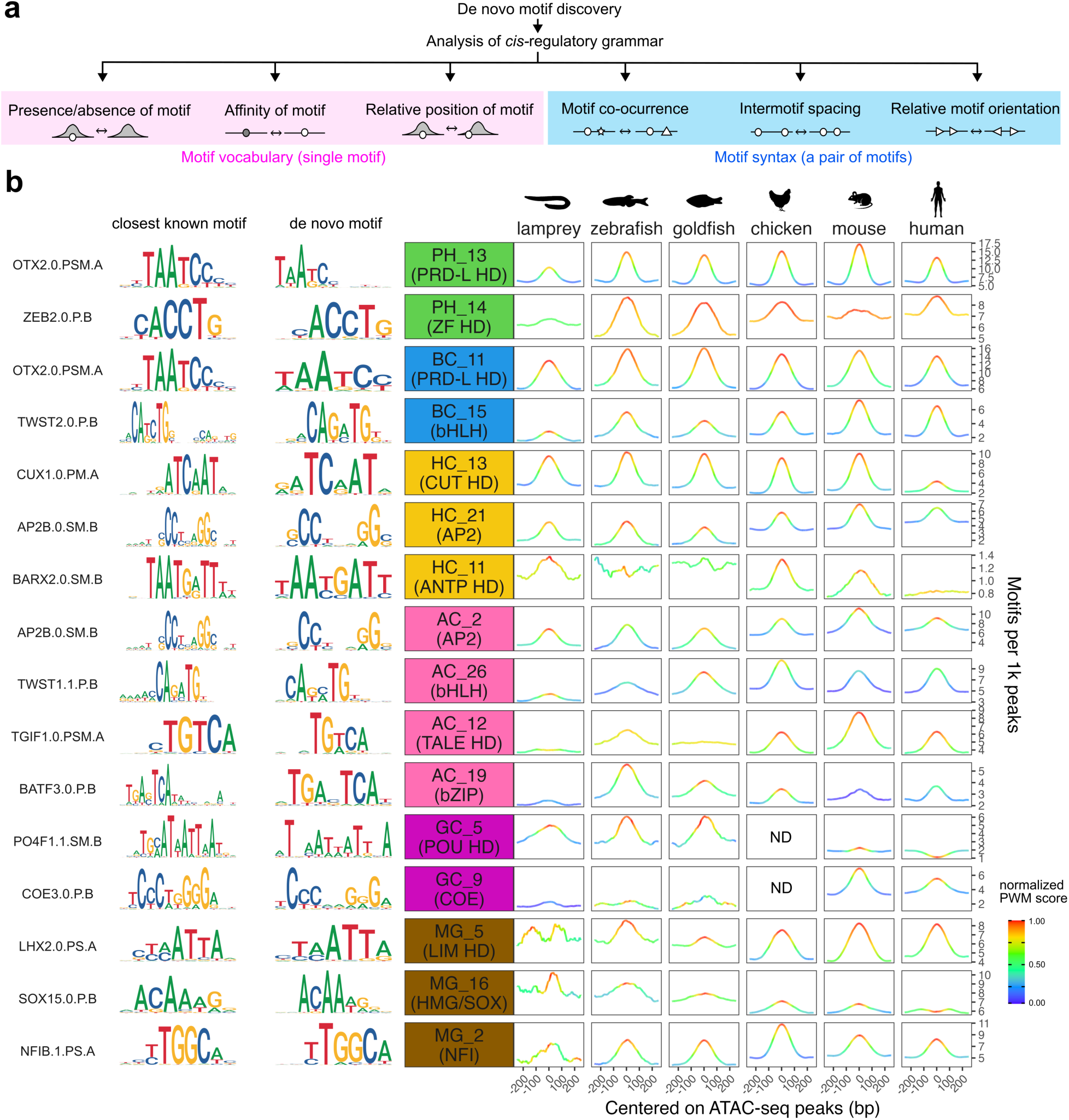
Evolutionary conservation of retinal *cis*-regulatory motif vocabulary distributions. **a,** Schematic representation of the major features of *cis-*regulatory grammar. **b,** The spatial distribution and normalized position weight matrix (PWM) scores of the EC motifs within class-enriched open chromatin regions in the six species are presented. The motifs are ordered from left to right according to motif enrichment as shown in Fig. 3d. The sequence logos for each EC motif and the closest known motif in the *HOCOMOCO* database^23^ are displayed on the left. Analysis of motif syntax is presented in **Supplementary Figs. 4 and 5**. PH, photoreceptor cell; BC, bipolar cell; HC, horizontal cell; AC, amacrine cell; GC, ganglion cell; MG, Müller glia; ND, not determined.

To ascertain whether syntactic features are conserved across species, we analyzed the co-occurrence, spacing, and relative orientation of all homo-and heterotypic pairs of EC motifs enriched in the same cell class (Figure 4a). As expected from the centralized pattern of enrichment of individual motifs (Fig. 4b), we observed enriched co-occurrence of all motif pairs—with the exception of GC_9 + GC_9—across all six species (Supplementary Fig. 4). In contrast, we observed little evidence for conserved patterns of relative motif spacing or orientation (Supplementary Fig. 5). One motif, BC_11, shows enrichment of tandem pairs with an intersite spacing of 8-11 bp (i.e., approximately one helical turn), but this pattern is only conserved across jawed species (i.e., not in lamprey). A similar pattern of co-occurrence of monomeric K50 homeodomain-type motifs was previously noted in CRX ChIP-seq peaks of mouse photoreceptors^29^. The related photoreceptor-enriched K50-type motif identified in the present study (PH_13) is a dimeric motif^18^. We therefore re-analyzed our photoreceptor-enriched OCRs using a monomeric K50 motif, which revealed helical co-occurrence of motif pairs similar to that observed for BC_11 (Supplementary Fig. 5b). Again, this co-occurrence pattern appears to be restricted to jawed species. We also detected a distinctive pattern of motif co-occurrence for the Müller glia-enriched HMG/SOX-type motif, MG_16, which consisted of pairs of motifs on opposite strands of the double helix, separated by four or five base pairs (Supplementary Fig. 5). This pattern of co-occurrence was conserved across all species including lamprey and likely represents a binding site for homo-or heterodimeric SOXE TFs (i.e., SOX8, SOX9, and SOX10), as previously described^30^. Indeed, we found that SOXE-type TFs showed Müller glia-enriched expression in all six species (Supplementary Table 4). We therefore consider this dimeric site to represent a single motif occurrence and not a feature of higher-order syntax. Thus, although almost all motif pairs show higher rates of co-occurrence than in control regions, few other higher-order syntactic features are shared between jawed and jawless species.

### Machine learning models of *cis*-regulatory grammar cluster by cell class

To enable quantitative comparison of class-specific *cis*-regulatory grammars across species, we trained machine learning models of grammar for each of the six cell classes in each of six species (with the exception of chicken ganglion cells). In light of the findings in the preceding section, we decided to build minimal vocabulary-based models that encompass only two key features of *cis*-regulatory grammar: the presence/absence of motifs and motif affinity. For this purpose, we constructed gapped *k*-mer support vector machine (gkm-SVM)^31^ classifiers to distinguish cell class-enriched OCRs (i.e., the positive training set) from broadly open chromatin regions (the negative training set). We constructed a total of 35 gkm-SVM models by randomly partitioning each training dataset into five subsets and performing 5-fold cross-validation (see Methods). We then used the five resultant models for each dataset to score the 35 test sets from each cell class and species. We measured the performance of the models by calculating the receiver operating characteristic (ROC) curve and the corresponding area under the curve (AUC) (Supplementary Fig. 6). For same-dataset validation, the mean ROC-AUC value for the 35 models was 0.854 (± 0.049 SD), with the best performance observed with the mouse horizontal cell model (0.920 ± 0.004 SD) and the worst performance with the lamprey amacrine cell model (0.733 ± 0.004 SD). Next, we evaluated the ability of the models to classify OCRs in the 34 other datasets. The average cross-species performance of models on other-class OCRs (e.g., lamprey photoreceptor model classifying mouse horizontal cell OCRs) was essentially random (ROC-AUC = 0.520 ± 0.086 SD) and provides an empirical estimate of baseline model performance. In contrast, for photoreceptor, bipolar cell, horizontal cell, and Müller glial models, we observed good performance in classifying same-class OCRs from different species, with all ROC-AUC scores above baseline performance, except for the lamprey Müller glial model whose performance was borderline overall, but consistently better for same-class OCRs than other-class OCRs (0.60-0.69 compared to 0.487 ± 0.030 SD) (Supplementary Fig. 6). The relatively poor performance of the lamprey Müller glial model is likely attributable to the small number of Müller glia identified in our snATAC-seq analysis and the corresponding paucity of class-enriched OCRs (151 sequences in total) available for model training (Supplementary Tables 5, 6). We observed comparable cross-species performance for amacrine and ganglion cell models, except for the zebrafish and goldfish models, which demonstrated excellent performance on each other’s datasets (ROC-AUC ≥ 0.80) but worse performance on non-teleost datasets (see Supplementary Fig. 6). Overall, the cross-species classificatory performance of these models confirms the existence of universally shared class-specific grammar features.

To evaluate the ability of these models to predict functionally important transcription factor binding sites across species, we used them to analyze the mouse *Gnb3* promoter, which drives expression in both photoreceptors and bipolar cells. We previously showed that this promoter contains five phylogenetically conserved K50 homeodomain binding sites, two of which are required for photoreceptor and bipolar expression^11^. We used photoreceptor and bipolar cell models trained on lamprey, zebrafish, goldfish, chicken, and human datasets to score the mouse *Gnb3* promoter. To visualize the contribution of individual nucleotides to the overall model scores, we used *GkmExplain*^32^, a feature attribution method which displays the predicted relative contribution of individual nucleotides as a sequence logo. All ten models (i.e., five photoreceptor and five bipolar cell models) produced highly concordant sequence logos, attributing particular importance to the two K50 motifs required for promoter activity (Supplementary Fig. 7). In fact, mutations of the highest-scoring nucleotides in all models (in motifs #2 and #4) result in a severe reduction of promoter activity (Supplementary Fig. 7). These findings demonstrate that *cis*-regulatory models from evolutionarily distant species are able to predict functionally important motifs—and even individual nucleotides—with great precision.

To evaluate the similarity and relatedness of models across species, we quantified the pairwise distances between models and used the resultant data to hierarchically cluster them. To accomplish this task, we first used the models to score all possible 11-mers, extracted the top 200 highest-scoring 11-mers for each model, and combined them to define a set of 4,276 unique 11-mers. We found that 53.8% (2,300 out of 4,276) of these 11-mers contain EC motifs, indicating that these models capture grammar features identified in the preceding section as well as additional features not detected by our motif-based approach. Next, we measured the pairwise distance between models by calculating the Pearson correlation coefficient between each model’s scores for the 4,276 11-mers (Fig. 5b). We then hierarchically clustered the models using one minus the Pearson correlation coefficient as a distance metric. The resulting hierarchical clustering revealed that the models group by cell class, not by species (Fig. 5c). The cell-class clusters aggregate into two super-clusters, one comprising photoreceptor and bipolar cell models, and the other comprising all other models. This higher-order grouping likely reflects the fundamental distinction between grammars dominated by the presence of K50 homeodomain binding site motifs (photoreceptor and bipolar cell grammars) and those that are not. Consistent with these results, we calculated a silhouette score—a metric used to evaluate the stability and robustness of clusters—and found that it culminated with the formation of six clusters (Supplementary Fig. 8). In summary, the robust co-clustering of retinal cell-class models from both jawed and jawless species highlights the deep evolutionary conservation of retinal *cis*-regulatory codes.

**Fig. 5.**
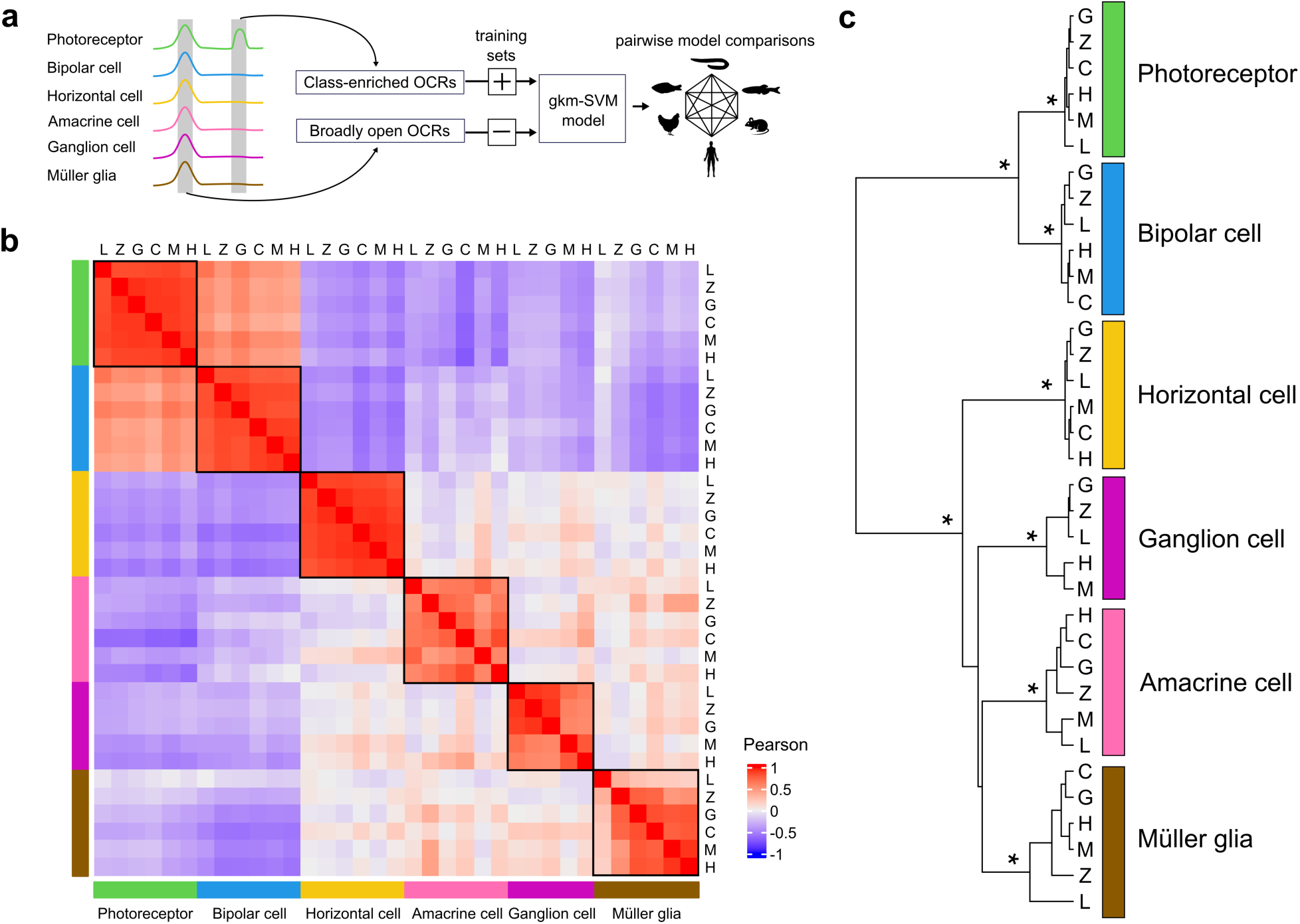
Retinal cell class-specific *cis*-regulatory grammars are conserved between jawed and jawless species. **a,** Schematic showing methodology used to generate and compare gkm-SVM models of *cis*-regulatory grammar. **b,** Heatmap showing Pearson correlation coefficients between gkm-SVM models. Models for a given cell class are enclosed by black boxes. (c) Hierarchical clustering of the gkm-SVM models. Branch nodes with a statistical significance of p < 0.01 are denoted by an asterisk (approximately unbiased p-value for selective inference by bootstrap resampling analysis followed by a multiscale resampling). L, lamprey; Z, zebrafish; G, goldfish; C, chicken; M, mouse; and H, human.

## Discussion

The fundamental cell class architecture of the vertebrate retina has remained largely unchanged over more than 500 million years of evolution. Utilizing single-cell chromatin profiles of retina from lamprey, zebrafish, goldfish, chicken, mouse, and human, we investigated *cis*-regulatory grammar and evaluated cross-species homology at the resolution of the cell class. Cross-species comparison of *cis*-regulatory grammar features revealed deep conservation of class-specific motif vocabulary but little preservation of higher-order syntax between jawed and jawless species. We identified between two and four EC motifs for each retinal cell class, underscoring the combinatorial nature of eukaryotic *cis*-regulatory codes^33^. Although we did not detect consistent patterns of motif spacing or orientation between jawed and jawless species, the central enrichment of motifs within OCRs results in a tendency for co-occurring motifs to cluster near the OCR summit. Pairwise comparison of machine-learning models of *cis*-regulatory grammar demonstrated close similarity of models within retinal cell classes, highlighting the evolutionary antiquity of vertebrate retinal *cis*-regulatory codes. We also observed higher-order grouping of models, possibly reflective of deeper ‘sister’ relationships among retinal cell classes as previously demonstrated for photoreceptors and bipolar cells^11^. Overall, these findings confirm that the six class-level *cis*-regulatory codes controlling vertebrate retinal gene expression arose in the common ancestor of extant vertebrates more than half a billion years ago and persist despite near-total enhancer replacement.

In the course of evolution novel cell types can arise via ‘duplication’ of a single ancestral cell type into two descendant daughter cell types, which subsequently evolve distinctive cellular features via a process known as ‘individuation’^2^. Shared cellular features may arise via evolutionary convergence in cell types derived from remote lineages.

Thus, the expression of effector genes controlling cell type-specific features—for example, the expression of opsins in photoreceptor types—is often a poor guide to the underlying evolutionary relationships of cell types. In contrast, the transcriptional regulatory networks which control expression of effector genes often persist for long evolutionary periods and are therefore more stable objects for evolutionary comparison. For this reason, Arendt and colleagues previously proposed that the presence of sets of terminal selector transcription factors—dubbed ‘core regulatory complexes (CoRCs)’—should be used to define cell types and trace their evolutionary origins^2,34^. The most common method for evaluating CoRCs is to measure the expression of TFs in individual cell types. However, most cell types express dozens of TFs, and it can therefore be difficult, without *a priori* knowledge, to prioritize factors for evolutionary comparison based on expression pattern alone. The present study introduces a methodologic solution to this problem: by elucidating the *cis*-regulatory codes of individual cell types or classes, it is possible to nominate the most likely cognate TFs that bind the enriched motifs which comprise the primary feature of *cis*-regulatory grammar.

Another reason why *cis*-regulatory codes may be a more reliable guide to deep evolutionary relationships than TF expression alone is that differential TF paralog choice may obfuscate shared patterns of TF usage in homologous cell types across species. For example, in mammals, the Maf family transcription factor *Nrl* is required for rod photoreceptor cell fate determination and gene expression^35^. Yet, while bird retinas contain rod photoreceptors^36^, avian genomes lack the *NRL* gene^37^. Instead, avian rods express another Maf transcription factor *MAFA*^38,39^, which is thought to play an analogous regulatory role to that of *Nrl* in mammals^39,40^. Similarly, while *nrl* is required for rod development in zebrafish larvae, it is dispensable for rod cell fate in adult fish^41^. This differential requirement might be explained by the fact that zebrafish co-express two Maf family members in rods, *nrl* and *mafba*, and either of these factors could contribute to maintenance of rod gene expression^42^. Thus, diverse vertebrates appear to have co-opted different Maf family TFs for the same regulatory purpose in rods. These various Maf family members bind similar motifs^43^, suggesting that shared patterns of motif enrichment within OCRs could inform evolutionary comparison despite differential paralog usage across species. Thus, in the face of both extensive enhancer turnover and differential TF paralog choice, comparison of *cis*-regulatory codes may be the most reliable method for defining interrelationships among evolutionarily distant cell types.

The remarkable evolutionary stability of *cis*-regulatory codes is likely attributable to their instantiation in thousands of enhancers across the genome and the lack of an evolutionary mechanism for coordinate changes in multiple genomic regions. If a mutation in a TF changes its DNA-binding preference, it could result in a catastrophic failure of binding across the genome and widespread perturbations of gene expression. Conversely, mutations in an individual enhancer motif might render the enhancer inactive unless its cognate TF undergoes simultaneous changes in its binding preference; yet such changes would, in turn, render the TF incapable of binding other enhancers. For these reasons, it appears that *cis*-regulatory codes and TF binding preferences are interlocked and exceptionally resistant to evolutionary change.

## Acknowledgements

We thank Mike White and Matt Toomey for providing useful comments on the manuscript. We thank the Genome Technology Access Core (GTAC) in the Department of Genetics at Washington University in St. Louis for library preparation and next-generation sequencing. We are grateful to Nicholas Johnson and the Hammond Bay Biological Station staff of the US Geological Survey for supplying lamprey. This work was supported by the National Institutes of Health (EY030075 to J.C.C.).

## Author contributions

Y.O. and J.C.C. conceived of the study and designed experiments. Y.L. generated the snATAC-seq library of the chicken retina and trained the gkm-SVM models. C.A.M. generated the scRNA-seq library for chicken retina and the Iso-Seq library of upstream-migrant lamprey. A.M. collected and dissected lamprey tissues under the supervision of G.L.F. and A.P.S. Y.O. performed all other experiments and computational analysis. Y.O. and J.C.C. prepared the manuscript and figures. J.C.C. supervised the project and obtained funding.

## Competing interests

The authors declare no competing interests.

## Methods

### Animal Husbandry

#### Lamprey

Downstream-and upstream-migrant sea lamprey (*Petromyzon marinus*) were provided to us by the Hammond Bay Biological Station of the US Geological Survey, Millersburg, MI, USA. Downstream migrant lamprey was captured by drift net in the St. Marys River while they were in the process of migrating to Lake Huron to begin the parasitic stage of the life cycle. Adult lamprey was captured in tributaries of Lake Huron (Ocqueoc River and Cheboygan River) in the process of their upstream spawning migration. Downstream-and upstream-migrant lamprey were kept in well-aerated tanks in cyclic 12L/12D h lighting in accordance with the rules and regulations of the NIH guidelines for research animals, as approved by the Institutional Animal Care and Use Committee of the University of California, Los Angeles.

#### Chicken

Fertilized specific-pathogen-free white leghorn chicken (*Gallus gallus*) eggs (Charles River Laboratories) were incubated at 38°C until hatching. Chicks were maintained in groups of 3–5 individuals under constant illumination from a heat lamp and were provided Purina Start & Grow medicated feed (Purina Animal Nutrition LLC) and tap water *ad libitum* until retinas were harvested. Chicken husbandry and experimental procedures were carried out in accordance with United States National Institutes of Health guidelines^44^ and approved by the Washington University in St. Louis Animal Care and Use Committee (protocol #22-0430).

### Lamprey retina RNA-seq and transcript annotation

To improve retinal gene annotations in lamprey, Iso-Seq was performed on retinas of upstream-migrants. Lampreys were deeply anesthetized with 400 mg/L tricaine methanesulfonate (MS-222; E10521, Sigma–Aldrich), decapitated, and enucleated. After removing the anterior chamber and the vitreous body from the eye, the retina was isolated from the retinal pigment epithelium in HEPES-buffered Ames’ solution at room temperature, flash frozen in liquid nitrogen, and stored at -80°C. Total RNA was extracted using Trizol and purified using an RNeasy Mini Kit (Qiagen). A SMRTbell Iso-Seq library was prepared according to the manufacturer’s protocol (Pacific Biosciences). The library was then sequenced using a single PacBio Sequel II SMRT cell. Demultiplexed PacBio circular consensus sequences (CCS)-generated HiFi reads with a predicted accuracy ≥Q20 were first processed using *lima* in (*isoseq3*; v.3.4.0; http://isoseq.how/) for 5’ and 3’ primer removal with parameters (--iso-seq --peek-guess). PolyA^+^ tails and artificial concatemers were trimmed and removed using *refine* (*isoseq3*) with the parameter (--require-polya), resulting in Full-Length Non-concatemer Reads (FLNC). Clustering was performed using the partial order alignment (POA) algorithm using *cluster* (*isoseq3*) with the parameter (--use-qvs). For the identification of unidentified isoforms in the lamprey retina, the resultant high-quality consensus sequences were mapped to the genome (kPetMar1) using *minimap2*^45^ with the following parameters: -ax splice -u f. *StringTie* (v2.2.1) was then used (with the command line option –L) to assemble a new transcriptome based on the mapped Iso-Seq reads. *StringTie* was then run again (with the command line option "merge -F 0 -T 0") to obtain an updated transcriptome annotation. This annotation incorporated modifications to transcript body definitions from the existing National Center for Biotechnology Information (NCBI) Reference Sequence (RefSeq) reference (GCF_010993605.1) and novel transcripts supported by the Iso-Seq reads.

### Gene orthology inference

To infer orthologous genes among evolutionarily diverse vertebrate species, we employed *Orthofinder*^28^, which defines an orthogroup (OG) as a set of orthologous and paralogous genes that can be used as an object for comparative analysis. In principle, an OG contains the set of genes that are descended from a single gene in the last common ancestor of all the species being considered. Several representative vertebrate and non-vertebrate species (vase tunicate, clubbed tunicate, hagfish, brook lamprey, skate, white shark, catshark, medaka, reedfish, gar, frog, coelacanth, green anole, platypus, and elephant) along with the six study species (lamprey, zebrafish, goldfish, chicken, mouse, and human) were used to infer OGs. OGs were defined by setting the ascidian node as a base node, and then hierarchical orthogroups (HOGs) were defined with a jawless-jawed vertebrate divergence node as a base node. For this purpose, protein sequences were obtained from the NCBI RefSeq or Ensembl databases, except for the lamprey and goldfish. For the latter species, the transcript sequences were retrieved from the genome with the transcript annotation using *gffread* (version 0.12.7), discarding multi-exon mRNAs that had any intron with a non-canonical splice-site consensus (i.e., not GT-AG, GC-AG, or AT-AC). Next, the sense strand of the gene transcripts was utilized to predict coding sequences via *TransDecoder* (version 5.5). This process employed the *TransDecoder.LongOrfs* function with the "-S" option. The longest sequence of predicted amino acids was selected for each gene and employed for orthology inference. The updated transcript assembly of the lamprey was employed, as described in the preceding section. Orthology inference was conducted using *Orthofinder* (version 2.5.5.2) with the following parameters: -M msa -S blast -A mafft -T fasttree. The species tree was subsequently corrected manually using the -s option. In lieu of the default *FastTree* tree inference program, *IQtree2* (version 2.3.2) was employed for the generation of all trees.

The resultant phylogenetic trees were subjected to a duplication-loss-coalescence analysis with the rooted gene trees to resolve speciation and gene duplication events. Finally, HOGs were identified by setting the jawless-jawed vertebrate divergence node as the root node. The nomenclature of genes in lamprey and goldfish was updated based on the information provided by HOGs. For example, two genes, "NTNG1-NTNG2-1" and "NTNG1-NTNG2-2", in lamprey are paralogs, and both are orthologs of the human genes NTNG1 and NTNG2. The accession numbers of gene annotations are provided in Supplementary Table 7.

### Sample collection and library preparation for single-cell sequencing

#### Lamprey

The retinas of downstream-migrant lamprey were dissected and snap-frozen as described above. Frozen nuclei were extracted using the Chromium Nuclei Isolation Kit (10x Genomics) according to manufacturer’s instructions. In brief, three retinas were dissociated in lysis buffer with a plastic pestle until a homogeneous solution was obtained. Residual tissue debris was removed with the Nuclei Isolation Column followed by centrifugation in Debris Removal Buffer. The supernatant was discarded, and the nuclei pellet was resuspended in Wash Buffer. The nuclei were pelleted by centrifugation and resuspended in 50 μl of Resuspension Buffer. A sample of nuclei was stained with propidium iodide and quantified using a hemocytometer. The remaining resuspended nuclei (∼18,000 nuclei per library) were utilized for transposition and loaded into the 10x Genomics Chromium Single Cell system. From a single nuclei suspension, two replicates of the multiome library were constructed using the Chromium Next GEM Single Cell Multiome Reagent Kit version 1 (10x Genomics), according to manufacturer’s instructions. The libraries were then subjected to sequencing on the Illumina NovaSeq platform.

#### Chicken

Newly hatched chicks were killed by carbon dioxide inhalation followed by manual cervical dislocation. Retinas were removed from the eye by dissection, transferred to calcium and magnesium-free Hank’s balanced salt solution (Thermo Fisher Scientific), and dissociated into single cells by papain digestion^11^. A single chick retina was incubated in 800 μl of HBSS with 1.3 mg of papain (Worthington Biochemical Corporation) at 37°C for 10 min. The retina was then dissociated by gentle trituration with a pipette. The dissociated cells were washed with Dulbecco’s Modified Eagle Media (DMEM; Thermo Fisher Scientific) containing 10% fetal bovine serum (Thermo Fisher Scientific) and DNaseI (Roche) for 5 min at 37°C. The cells were then pelleted by centrifugation at 1000 × *g* for one minute. The cells were resuspended in Dulbecco’s phosphate-buffered saline (DPBS) with 1% BSA and filtered. To isolate nuclei, dissociated cells were subjected to centrifugation at 300 × *g* for 5 min at 4°C. The cell pellet was then resuspended in 100 µl of lysis buffer [0.1% IGEPAL CA-630, 0.1% Tween-20, and 0.01% digitonin in lysis dilution buffer (10 mM Tris-HCl, 10 mM NaCl, 3 mM MgCl2, 1% BSA, pH 7.4)], mixed three times by pipetting, and incubated on ice for 3 min. The nuclei were then added to 1 ml of wash buffer (0.1% Tween-20 in the lysis dilution buffer) and centrifuged at 500 × *g* for 5 min at 4°C. Finally, the nuclei pellet was resuspended in 50 µL of 1x Diluted Nuclei Buffer (10x Genomics) and filtered through a 40 µm Flowmi™ Cell Strainer. The nuclei were quantified using a hemocytometer. ∼22,000 nuclei were then resuspended and used for transposition and subsequently loaded into the 10x Genomics Chromium Single Cell system. The scATAC-seq library was constructed using the Chromium Single Cell ATAC Reagent Kits v1.1 (10x Genomics), according to manufacturer’s instructions. The libraries were subjected to sequencing on the Illumina NovaSeq platform.

### Sequencing data preprocessing

Publicly available data were retrieved from previous studies, including snATAC-seq datasets for goldfish^46^ and mouse^47^ as well as multiome (snRNA-seq + snATAC-seq) datasets for zebrafish^48^ and human^49^. The processed sequencing data was retrieved from the original study in zebrafish, while for the other five species, the raw sequence data were initially processed by the Cell Ranger ATAC version 2.0.0 pipeline or the Cell Ranger ARC version 2.0.0 pipeline (10x Genomics) for read filtering, alignment against the genome, and barcode counting. The genome assemblies used were kPetMar1 (lamprey), GCA_014332655.1 (goldfish), galGal6 (chicken), mm10 (mouse), and hg38 (human). The processed data were loaded into *ArchR*^50^ (version 1.0.2) for goldfish, chicken, and mouse, while the lamprey and human multiome data were loaded into *Seurat*^51,52^ (version 4.0.3) and *Signac*^53^ (version 1.3.0). Transcriptome annotations used in this study are as follows: lamprey, custom annotation as described in the previous section; goldfish, the previously described custom annotation^46^; chicken, NCBI *Gallus gallus* Annotation Release 104; mouse, mm10-2020-A-2.0.0 provided by 10x Genomics; and human, GRCh38-2020-A-2.0.0 provided by 10x Genomics.

### snATAC-seq analysis

#### Lamprey

Multiome (snRNA-seq + snATAC-seq) sequence reads from upstream-migrant lamprey retina were used as input data for analysis by *Seurat* and *Signac*, respectively. Low-quality cells were removed if the cell contained <300 expressed genes; if >0.3% of the cell’s total gene expression derived from mitochondrial genes; if <1,000 fragments were detected; or if the cell had a transcription start site enrichment score <3. Putative doublets were removed based on the presence of >40,000 fragments or >10,000 expressed genes.

For gene expression analysis, the negative binomial regression normalization method implemented in *SCTransform* (*Seurat*) was used to standardize the gene expression matrix and reduce gene expression noise. Dimensionality reduction of gene expression was then conducted with *RunPCA*. The significance of each principle component in the RNA analysis was evaluated manually using elbow plots. 1 to 30 dimensions were chosen for the subsequent analysis.

A cell-by-region count matrix of ATAC-seq reads overlapping open chromatin regions defined by *Cell Ranger* ATAC was generated using *FeatureMatrix* and *CreateChromatinAssay* (*Signac*). The count matrix was subjected to normalization and dimensionality reduction using *FindTopFeatures* (*Signac*) with a min.cutoff of ‘q0’, *RunTFIDF* (*Signac*), and *RunSVD* (*Signac*) with default settings. The correlation between total counts and each dimension in the ATAC analysis was visualized with *DepthCor* (*Signac*), and the first Latent Semantic Indexing (LSI) component, which captured sequencing depth (technical variation), was filtered.

The nearest neighbors for each cell were identified based on a weighted combination of two modalities (RNA and ATAC) by constructing a weighted nearest neighbor (WNN) graph using *FindMultiModalNeighbors* with 1 to 30 dimensions (PCA) and 2 to 30 dimensions (LSI). The clusters were determined by modularity optimization using *FindClusters* with a resolution of 0.2.

Cell clusters were assigned to retinal cell classes based on the expression of the following cell-class marker genes^9^: *RHO* and *GNAT2* (photoreceptor), *SLC17A6_1, SLC17A6_2, GRIK2_1, GRIK2_2,* and *PRKCA* (bipolar cell), *ONECUT1* (horizontal cell), *SLC6A9, SLC6A11_1, SLC6A11_2,* and *SLC32A1* (amacrine cell), and *RBPMS* (ganglion cell). The Müller glia cluster was manually annotated using *CellSelector* (*Seurat*) based on the expression of *GLUL*.

Two technical replicates were independently analyzed. Following cell annotation in each of the two technical replicates, the data were merged into a single dataset. Gene expression data were then normalized, scaled, and subjected to PCA analysis as described above. The clusters were identified by constructing a *k*-nearest neighbor graph with *FindNeighbors* (*Seurat*) with 1 to 40 dimensions (PCA), followed by modularity optimization using *FindClusters* (*Seurat*) with a resolution of 1.5. The cell embedding was obtained with *RunUMAP* (*Seurat*). The cell annotations were transferred from the multimodal clustering analysis conducted in each sample. Finally, putative mixed clusters (i.e., those assigned annotations for more than one cell class) were removed from the analysis.

#### Zebrafish

Multiome (snRNA-seq + snATAC-seq) sequence reads from adult zebrafish retina were retrieved from a repository (see original study^48^) and used as input data for analysis by *Seurat* and *Signac*, respectively. Low-quality cells were removed if > 5% of the cell’s total gene expression derived from mitochondrial genes; if < 1,000 fragments were detected; or if the cell had a transcription start site enrichment score <2. Putative doublets were removed based on the presence of >6,000 fragments or > 4,000 expressed genes. Cell annotations presented in the original study^48^ were used.

#### Goldfish

Barcoded and aligned fragments were used as input data for ATAC analysis by *ArchR*. A genome-wide tile matrix with insertion counts was calculated on 500-bp non-overlapping windows using *createArrowFiles*. Low-quality cells were removed if they had a transcription start site enrichment score <15 or <1,000 fragments. Putative doublets were removed using *filterDoublets* with a filterRatio of 2. Nuclei with >50,000 fragments were also excluded.

Dimensionality reduction was implemented with *addIterativeLSI* using the LSI method. Cell clustering was then performed with *addClusters* using a shared nearest neighbor (SNN) modularity optimization-based clustering algorithm. A total of eight clusters were identified, comprising rod photoreceptors, cone photoreceptors, bipolar cells, horizontal cells, amacrine cells/ganglion cells, Müller glial cells, microglia, and oligodendrocytes. Cluster annotation was performed based on marker genes identified in the original study^46^. To distinguish between amacrine and ganglion cells, cell annotations from scRNA-seq data were projected onto cells in the snATAC-seq dataset. First, 501-bp open chromatin regions were generated with *addReproduciblePeakSet*, utilizing the pseudo-bulk ATAC replicates for each cluster. Next, peak sets from each cell class were merged to create a non-redundant union set, which was then subjected to analysis in *Signac* for normalization and dimensionality reduction, as described above for lamprey. Clusters were identified using *FindClusters* with a resolution of 2.0. The processed data were then integrated with the scRNA-seq data, where the gene expression profile was processed using a standard protocol in *Seurat* and the cell clusters were annotated according to the expression of marker genes identified in the original study^46^. Cell annotations for the scRNA-seq data were then projected onto the cells of the snATAC-seq dataset using *FindTransferAnchors* with the cca reduction method, followed by *TransferData* with the LSI weight reduction method, using 2 to 30 dimensions. Transferred labels were retained based on the most abundant cells within each cluster. Only the five classes of retinal neurons and Müller glia were retained for subsequent analysis.

#### Chicken

Barcoded and aligned fragments were used as input data for ATAC analysis by *ArchR*. Low-quality nuclei were removed if they had a transcription start site enrichment score <15 or <3,000 fragments. Putative doublets were removed using the *filterDoublets* function with a filterRatio of 3. Nuclei with >25,000 fragments were also excluded.

Dimensionality reduction was implemented by the LSI method using *addIterativeLSI* with two iterations and cluster parameters of resolution of 0.1. Cell clustering was then performed using a shared nearest neighbor (SNN) modularity optimization-based clustering algorithm, employing *addClusters* with a resolution of 0.4. Retinal cell classes were identified by enrichment of the following marker genes^54^: *GNAT2* and *RHO* (photoreceptors), *VSX1* and *VSX2* (bipolar cells), *ONECUT3* (horizontal cells), *SLC32A1* (amacrine cells), *RLBP1* (Müller glia), and *OLIG2* (oligodendrocytes). To verify that ganglion cells were absent from our dataset, we projected cell annotations for published scRNA-seq data^54^ onto the cells in the snATAC-seq data, as described above for goldfish. We failed to detect any ganglion cell clusters. Only the four other classes of retinal neurons and Müller glia were retained for subsequent analysis.

#### Mouse

Barcoded and aligned fragments from multiple developmental stages (E11, E12, E14, E16, E18, P0, P5, P8, P11, and P14) were pooled and used as input data for analysis by *ArchR*. Low-quality cells were removed if they had a transcription start site enrichment score <10. Putative doublets were removed using the *filterDoublets* function with a filterRatio of 2. Nuclei with >30,000 fragments were also excluded.

Dimensionality reduction and cell clustering was performed using *addClusters* with a resolution of 0.7. A total of fourteen clusters were identified, comprising rod photoreceptors, cone photoreceptors, bipolar cells, horizontal cells/amacrine cells, ganglion cells, Müller glial cells, early cone photoreceptors, early rod photoreceptors, early neurogenic cells, late neurogenic cells, early retinal progenitor cells, and three stages of retinal progenitor cell. Cluster annotation was performed based on marker genes identified in the original study^47^. To distinguish between horizontal cells and amacrine cells, we projected cell annotations for developmental-stage-matched scRNA-seq data^55^ onto the cells in the snATAC-seq data, as described above for goldfish, except for using *FindClusters* with a resolution of 0.4. Only mature retinal neurons and Müller glia were retained for subsequent analysis.

#### Human

Multiome (snRNA-seq + snATAC-seq) data were processed in a manner analogous to that described in the original study^49^. In brief, for each sample, count matrices for both RNA-seq and ATAC-seq were loaded into *ArchR*. Low-quality cells were removed if they had <200 RNA transcripts, >0.8% mitochondrial gene transcripts, <3,000 ATAC fragments, or a transcription start site enrichment score <7. Putative doublets were removed using the *filterDoublets* function with a filter ratio of 5. Additional doublets we removed if cells expressed >25,000 RNA transcripts, >7,000 genes, or >70,000 ATAC fragments. The remaining nuclei in all preprocessed samples were subsequently merged into a single *Seurat* object. Gene expression counts were normalized using *NormalizeData*, scaled using *ScaleData*, and batch-corrected using *Harmony*^56^. Graph-based clustering was then performed on the *Harmony*-corrected data using the top 20 principal components at a resolution of 0.5. Cluster annotation was performed based on marker genes identified in the original study^49^. Clusters co-expressing marker genes from different cell classes were excluded; clusters devoid of any marker genes were also excluded. Only the five classes of retinal neurons and Müller glia were retained for subsequent analysis.

### scRNA-seq analysis

Single-cell RNA-seq data for lamprey, zebrafish, and human derive from multiome datasets described in the preceding section. For mouse, scRNA-seq data and corresponding cell cluster annotations were retrieved from a single-cell expression atlas of adult retina^57^. For goldfish, the raw sequence data were obtained from a published study^46^ and processed using *Cell Ranger* (version 7.1.0; 10x Genomics) for read filtering, alignment against the genome, and barcode counting. Goldfish retinal cell-class annotations were also retrieved from the paper^46^, and cells that were both retained in our analysis and included in their original cell annotations, were used for subsequent analysis. For chicken, single-cell RNA sequencing data was generated in the present study from newly hatched (P0) chicken retinas. The raw sequence data was processed using *Cell Ranger* (version 7.0.0), and the aligned gene expression reads were loaded into *Seurat*. Low-quality cells were removed if the cell contained <1,000 expressed genes or if >10% of the cell’s total gene expression derived from mitochondrial genes. Putative doublets were removed based on the presence of >15,000 expressed genes. Count data were normalized using *SCTransform* (*Seurat*), and dimensionality reduction was performed using *RunPCA* (*Seurat*). The significance of each principle component in the RNA analysis was evaluated manually using elbow plots. 1 to 40 dimensions (PCA) were used for identifying the clusters with *FindNeighbors*, followed by modularity optimization with *FindClusters* using a resolution of 1.2. Retinal cell classes were identified based on the expression of cell-class marker genes^54^. Raw data acquisition and processing will be presented in a forthcoming publication.

### Cell-class meta-gene analysis

To devise a metric for measuring evolutionarily conserved cell-class meta-gene activity, differentially expressed genes were first identified using the Wilcoxon rank sum test. Single-cell RNA-seq datasets from the six study species (described above) were analyzed using *FindAllMarkers* (*Seurat*) with the following parameters: logic.threshold = 0, min.pct = 0.05, return.thresh = 1. Differentially expressed genes were defined as those which exhibited an average log_2_-fold change > 0.1, an adjusted p-value < 0.01, and a percentage of cells expressing the gene >10%. Differentially expressed genes were further filtered for cell-class-enriched expression as follows. Pseudo-bulk gene expression was quantified for each cell class using *AverageExpression* (*Seurat*), and then tau (τ), a measure of cell-class specificity of gene expression, was calculated^58^. Tau 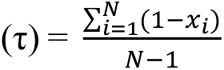, where *N* is the number of cell classes, and *x_i_* is the expression profile component normalized by the maximal expression value. Genes with τ > 0.6 were retained for subsequent analysis.

The top 1,000 genes with the highest tau index were identified for each cell class in each species and then intersected across the six study species to identify evolutionarily conserved differentially expressed genes. To accomplish this task, hierarchical orthogroups (HOGs) were defined using *Orthofinder* described above in **Gene orthology inference**. HOGs that contained differentially expressed genes from lamprey and three or more jawed species were retained, and for each species, the cell-class-enriched genes included in the HOGs were defined as evolutionarily conserved differentially expressed (ecDE) genes. Next, chromatin openness over the promoter and gene body of ecDE genes was quantified using *GeneActivity* (*Signac*). Individual gene activity scores were aggregated into a single activity score, referred to as ‘cell-class meta-gene activity’. Meta-gene activity was quantified in individual cells for each cell class, and the gene accessibilities of both the meta-genes and the remaining genes were normalized and scaled in accordance with the standard protocol in *Signac*. The statistical significance of cell-class enrichment of the meta-genes was determined by comparing the cell class exhibiting the highest score for the meta-gene with the second-highest scoring cell class using the Wilcoxon rank sum test followed by Bonferroni correction, performed using *Wilcox.exact* in the *exactRankTests* package in *R*.

### Identifying differentially accessible open chromatin regions

Two replicates of pseudo-bulk ATAC data were generated from subsets of snATAC-seq data for each retinal cell class using *addGroupCoverages* (*ArchR*). 201-bp consensus open chromatin regions (peaks) were generated using *addReproduciblePeakSet* for the replicates for each cell class with the following parameters: extendSummits = 100 and cutOff = 0.1. The peak sets from each cell class were then combined, and the resultant non-redundant union peak set was used for the following analysis. Single-cell differential accessibility tests were performed using *FindAllMarkers* (*Seurat*) with the following parameters: logfc.threshold = 0, only.pos = TRUE, min.pct = 0.01, test.use = LR, latent.vars = nCount_ATAC, return.thresh = 0.01, slot = counts. Peaks with an adjusted p-value <0.01 and an average log_2_-fold change >0 were retained. Additionally, peaks were retained if their average accessibility in pseudo-bulk ATAC-seq data— calculated with *AverageExpression* (*Seurat*)—was four times greater than the average of the average accessibilities from the other cell classes. Furthermore, cell-class-enriched peaks were filtered if they did not overlap with peaks called in the corresponding cell type. The resultant peak sets were defined as differentially accessible peaks. Broadly open peaks were defined as all peaks not contained in the union of differentially accessible peaks for each species. Chromatin accessibility of cell-class-enriched open chromatin regions was visualized using *deepTools*^59^ (version 3.5.4). The number of OCRs used for the following analysis is provided in Supplementary Table 6.

### Determining cross-species alignability of cell-class-enriched OCRs

The union of all differentially accessible OCRs for each of five study species (lamprey, zebrafish, chicken, mouse, and human) was mapped onto the reference genomes of various vertebrate species. Regions that overlapped with exons were excluded prior to mapping. The UCSC Genome Browser’s *LiftOver* utility^60^ (version 377 in bioconda) was employed for mapping, utilizing a parameter minMatch=0.5. For this analysis, precomputed reciprocal best-hit whole-genome alignment (rbest.chain) files were downloaded from the UCSC Genome Browser website. Differentially accessible OCRs in zebrafish were mapped from danRer11 onto danRer7 and then mapped onto the reference genomes of other species, utilizing the publicly available reciprocal best chain files. Mapping between conspecific reference genomes was conducted using the *LiftOver* utility with a parameter minMatch=0.95 and the “over.chain” file. Similarly, lamprey OCRs were mapped from kPetMar1 onto petMar3 with the custom chain file generated using *flo*^61^, a UCSC Genome Browser command line wrapper. The evolutionary divergence times of all internal branches were obtained from *timetree*^62^.

The decay of sequence mappability (i.e., alignability) was modeled using a fitted Gompertz equation 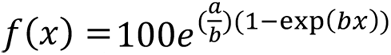, with divergence time as a variable. Equation fitting was performed using the Gauss-Newton algorithm with starting estimates (a = 0.001, b = 0.01) and a maximum of 1,000 iterations allowed in the *nls* function in the *stats* package (*R*; version 4.1.0). The resultant fitted parameter values were a = 0.002508772 and b = 0.006060751. The confidence intervals were determined by 10,000 bootstrap samples using the *boot_nls* function in *nlraa* (version 0.89, https://github.com/femiguez/nlraa).

### Discovery of evolutionarily conserved motifs

#### De novo motif discovery

*De novo* motifs (position probability matrices) were identified for each species using *findMotifsGenome.pl* (HOMER v4.11)^21^, with parameter -size 201. Cell-class-enriched OCRs were used as target sequences, and the broadly open regions were used as background sequences for each species. Identified motifs were retained for subsequent analysis if the statistical significance of the motif was <10^-10^ and if the motif was present in >2% of the cell-class-enriched OCRs.

#### Motif Clustering

The motif clustering approach was adapted from a prior study^63^. For each cell class, *de novo* motifs of all study species were subjected to pairwise comparison using *Tomtom* (MEME Suite; version 5.4.1)^64^, with the following parameters: -dist kullback, -motif-pseudo 0.1, -min-overlap 1. The pairwise comparison similarity values (E-values) were calculated by multiplying each *Tomtom*-reported p-value by the number of target motifs, and a group of -log_10_(E-values) for each motif was then employed to measure the Pearson correlation coefficient between two motifs. E-values that exceeded 100 were capped at a maximum value of 100. Hierarchical clustering was performed using the average linkage comparison method with one minus the Pearson correlation coefficient as the distance metric. *hclust* in the *stats* package (*R*; version 4.1.0) was used for clustering. Motif clusters were identified by cutting the resultant dendrogram at a height of 0.9. To eliminate clusters with dissimilar motifs, the median value of the Pearson correlation coefficient among the most significantly enriched motifs from each species within a cluster was calculated, and those clusters with median values ≤0.5 were removed. Motif clusters were defined as ‘evolutionarily conserved’ if they included motifs from lamprey and three or more jawed species. Given the absence of chicken ganglion cells in our dataset, evolutionary conserved motif clusters in ganglion cells were only required to contain motifs from lamprey and two or more jawed species.

To create consensus ‘merged’ motifs, the most significantly enriched motifs from each species in each retained cluster were subjected to sequence alignment, and for each position, the mean value of the position probability matrix was calculated using *mergeMotifs* (*motifStack*^65^, version 1.38.0). Undefined flanking regions for each motif were assigned a probability of zero and included in the mean calculation. The merged motifs were converted from a position probability matrix to an information content matrix, and then the motif flanking region was trimmed from either side if its information content was <0.05. The resultant merged motifs were then compared with mammalian TF binding motifs in the *HOCOMOCO* database^23^ using *Tomtom*^22^.

### Analysis of *cis-*regulatory grammar

#### Motif scanning

To identify occurrences of motifs, OCRs were scanned for motifs using *scan_sequences* (*universalmotif*; version 1.16.0; https://github.com/bjmt/universalmotif), retaining the highest-scoring motif if two motifs overlapped. The cutoff thresholds for calling motifs were calculated as the median of thresholds identified by HOMER across species. The background nucleotide frequencies were set to those observed in the corresponding cell-class-enriched 201-bp OCRs for each of the six species.

#### Motif distribution and affinity

To graph motif distributions and affinities within OCRs, the median PWM score and motif density per nucleotide were calculated. The per-nucleotide values were further smoothed using a 101-bp sliding window centered on the nucleotide in question.

#### Motif co-occurrence

For each pair of motifs, the number of OCRs with at least one pair was counted. Co-occurrences involving two overlapping motifs, regardless of the strand, were excluded. The enrichment of co-occurrence was then calculated as the ratio of the frequency of motif pairs in cell-class-enriched OCRs to the frequency of motif pairs in background OCRs. 50,000 background sequences were semi-randomly selected from the set of broadly open regions using *homer2 bg* (*HOMER*; version 5.1) with parameters: -ikmer 2 -N 50000 -NN 100000000. In this way, the distribution of dinucleotide frequencies in background sequences was matched to that in the target cell-class-enriched OCRs.

#### Motif spacing

To identify preferential spacing between motifs, the distance between the primary and secondary motifs in cell-class-enriched OCRs was quantified. Two relative distances to the primary motif were measured: the distance between the 5’ nucleotide of the primary motif and the 3’ nucleotide of the secondary motif, and the distance between the 3’ nucleotide of the primary motif and the 5’ nucleotide of the secondary motif. The distance between the two motifs was determined by selecting the value closest to zero. If the secondary motif was found to be 3’ downstream of the primary sequence, the length of the primary motif was added to this distance, ensuring that the distance was always measured from the 5’ nucleotide of the primary motif. The number of instances of the secondary motif at all nucleotide positions on both strands between ±40 bp relative to the primary motif was tallied.

### Nominating cognate transcription factors for evolutionarily conserved motifs

To identify cognate TFs that might bind evolutionarily conserved motifs, merged motifs were compared with mammalian TF binding motifs in the HOCOMOCO database^23^ using *Tomtom*. To nominate cognate TFs across the six study species, we assumed that homologous TFs belonging to the same hierarchical orthologous group (HOG; see **Gene orthology inference**) exhibit similar binding preferences. Zinc finger proteins belonging to the largest orthogroup were excluded from this analysis on account of the extensive divergence of binding preferences among zinc finger proteins, which is well-attested^66^.

Next, single-cell RNA-seq data were employed to identify TFs exhibiting cell-class-enriched expression in each of the six study species. Differential expression was determined using the Wilcoxon rank sum test in *FindAllMarkers* (*Seurat*). Pseudo-bulk expression profiles of TF genes in each cell class of each species were quantified using *AverageExpression* (*Seurat*) to assess the degree of cell-class enrichment for each TF. Candidate cognate TFs were retained if the corresponding motif similarity value (q-value in the *Tomtom* output) was <10^-1^ and the significance of the cell-class enrichment (adjusted p-value in the output of the differential expression test) was <10^-1.5^.

### Training and comparison of gkm-SVM models

#### Model training and validation

201-bp OCRs identified from snATAC-seq data in the six study species were used for training gkm-SVM models^31^. Differentially accessible OCRs in each of the six retinal cell classes were utilized as the positive training set, and broadly open OCRs from the same species were used as the negative training set. For each species, the same number of positive and negative OCRs were used for training and testing. The datasets were randomly divided into five equal groups, ensuring that each group was non-overlapping, and a 5-fold cross-validation was performed. Models were trained using *LS-GKM* ^67^ (version 0.1.0) with the following parameters: a linear gkm kernel without center weight (kernel 2), L=11, K=7, C=1.

Trained models were utilized to score all test sets within each cell class of each species. ROC-AUC scores were calculated based on the scores of the corresponding positive and negative test sets using *ROCR*^68^ (version 1.0-11). The overall performance of the models was determined by measuring the mean and standard deviation of ROC-AUC scores from the five independent replicates of the 5-fold cross-validation models.

#### Model interpretation

The contribution score of each nucleotide in the OCRs to the classification was computed using *gkmexplain*^32^ (release version 1.0.0). The importance scores for the mouse *Gnb3* promoter were calculated for photoreceptor and bipolar cell models of all study species except mouse. Both importance and hypothetical importance scores were calculated using the *gkmexplain* command, and the importance scores were normalized to the hypothetical importance score per developer’s recommendation. The resultant normalized importance scores were averaged among five model replicates obtained by 5-fold cross-validation (described above).

#### Model clustering

All possible 11-mer sequences were scored with the gkm-SVM models using the *gkmpredict* command. The resultant scores were averaged among five model replicates obtained by 5-fold cross-validation for each cell class of each species. The 200 highest-scoring 11-mers were selected for each model and merged to create a union of the best-scoring 11-mers (4,276 11-mers in total). Agglomerative hierarchical clustering was performed using one minus the Pearson correlation coefficient as a distance metric and Ward’s minimum variance method, employing *hclust* in the *stats* package (*R*; version 4.1.0). The statistical significance of a branching node was calculated using the *pvclust* package (version 2.2-0, https://github.com/shimo-lab/pvclust), where the approximately unbiased p-value for selective inference (SI p-value) was calculated by bootstrap resampling analysis followed by a multiscale resampling implemented in *pvclust*. The tree was visualized using the *ggtree* package (version 3.10.1). The silhouette score of each gkm-SVM model was measured using *silhouette* in the *cluster* package (version 2.1.2). The resulting scores were averaged across models within a cluster and subsequently averaged across clusters to evaluate cluster robustness.

### Code and data availability

All raw single-cell sequencing data and processed data generated in this study are available through the Gene Expression Omnibus (accession number pending). Cell metadata for scRNA-seq and snATAC-seq, hierarchical orthogroups, genomic coordinates of open chromatin regions, evolutionarily conserved motif PWMs, and gkm-SVM models are available in FigShare (accession number pending). The code used for the analysis is available upon request.

## Supplementary Tables

**Supplementary Table 1 | Cross-species sequence alignability of retinal cell class-enriched open chromatin regions.** This table includes the data used to create Fig. 2. All column headers are presented in italics. *Divergence_time*, divergence time between query and target species in *Mys*, millions of years; *Percentage_alignable,* the percentage of query sequences alignable to the target genome assembly; *Gompertz_prediction*, the value of the fitted Gompertz equation at the indicated divergence time; *Q2.5* and *Q97*, the boundary values of the 95% confidence interval as estimated for the fitted Gompertz equation at the indicated divergence time by bootstrap resampling (see Methods).

**Supplementary Table 2 | Gene expression profiles of evolutionarily conserved marker genes used to determine meta-gene accessibility.** This table includes the results of three analyses: the differential gene expression test performing using the Wilcoxon rank sum test on scRNA-seq data (*Cell_class*, *Ave_log_2_FC*, and *p_val_adj*); gene orthology inference (*HOG*); and the cell-class expression specificity index [*tau (τ)*]. See Methods for further details.

**Supplementary Table 3 | Enriched *de novo* motifs discovered by *HOMER*.** All *de novo* motifs found by *HOMER* to be enriched in cell class-enriched open chromatin regions of all six species are present in six sheets, one for each cell class. *ClusterID*, the identifier of the cluster to which this motif belongs as determined by hierarchical clustering; *Rank*, the rank order of the statistical significance of enrichment of this motif among all motifs in this cell class and species; Sequence, sequence of *de novo* motif; *PWM_threshold*, the PWM log-odds score threshold for calling the motif; *PosN*, the number of motif hits in target open chromatin regions; *PosT*, the total number of target open chromatin regions; *NegN*, the number of motif hits in background control sequences (non-integer values are present in the *NegN* column because *HOMER* employs weights for each background control sequence to normalize sequence bias between target and background open chromatin regions); *NegT*, the total number of background control sequences; *Enrichment*, motif enrichment =(*PosN/PosT*)/(*NegN/NegT*); *log_10_*, the statistical significance of motif enrichment measured using a binominal distribution. The probability of either of two possible outcomes (i.e., presence or absence of the motif in a region) was defined by *NegN/NegT* (motif presence) and 1 – *NegN/NegT* (motif absence); *Z-score*, the z-score in the binomial distribution.

**Supplementary Table 4 | Assignment of evolutionarily conserved motifs to cognate transcription factors.** *EC_motif*, evolutionarily conserved motif; *TF_motif*, identifier of the motif in the *HOCOMOCO* database that most closely resembles the EC motif; *TF_family*, the transcription factor family to which the TF that binds *TF_motif* belongs; *Motif_similarity*, the minimal false discovery rate at which the observed similarity would be deemed significant (i.e., the q-value in the *Tomtom* output); *Cell_class*, the cell class in which the gene is significantly enriched relative to other cell classes; *p_val_adj*, Bonferroni-corrected p-value determined by the differential gene expression test using the Wilcoxon rank sum test on scRNA-seq data; *HOG*, hierarchical ortholog group.

**Supplementary Table 5 | The number of cells of each retinal cell class in snATAC-seq datasets of the six species**

**Supplementary Table 6 | The number of open chromatin regions used for *de novo* motif discovery and gkm-SVM model training**

**Supplementary Table 7 | Accession numbers of sequences used for orthology inference**

---

**Supplementary Fig. 1.**
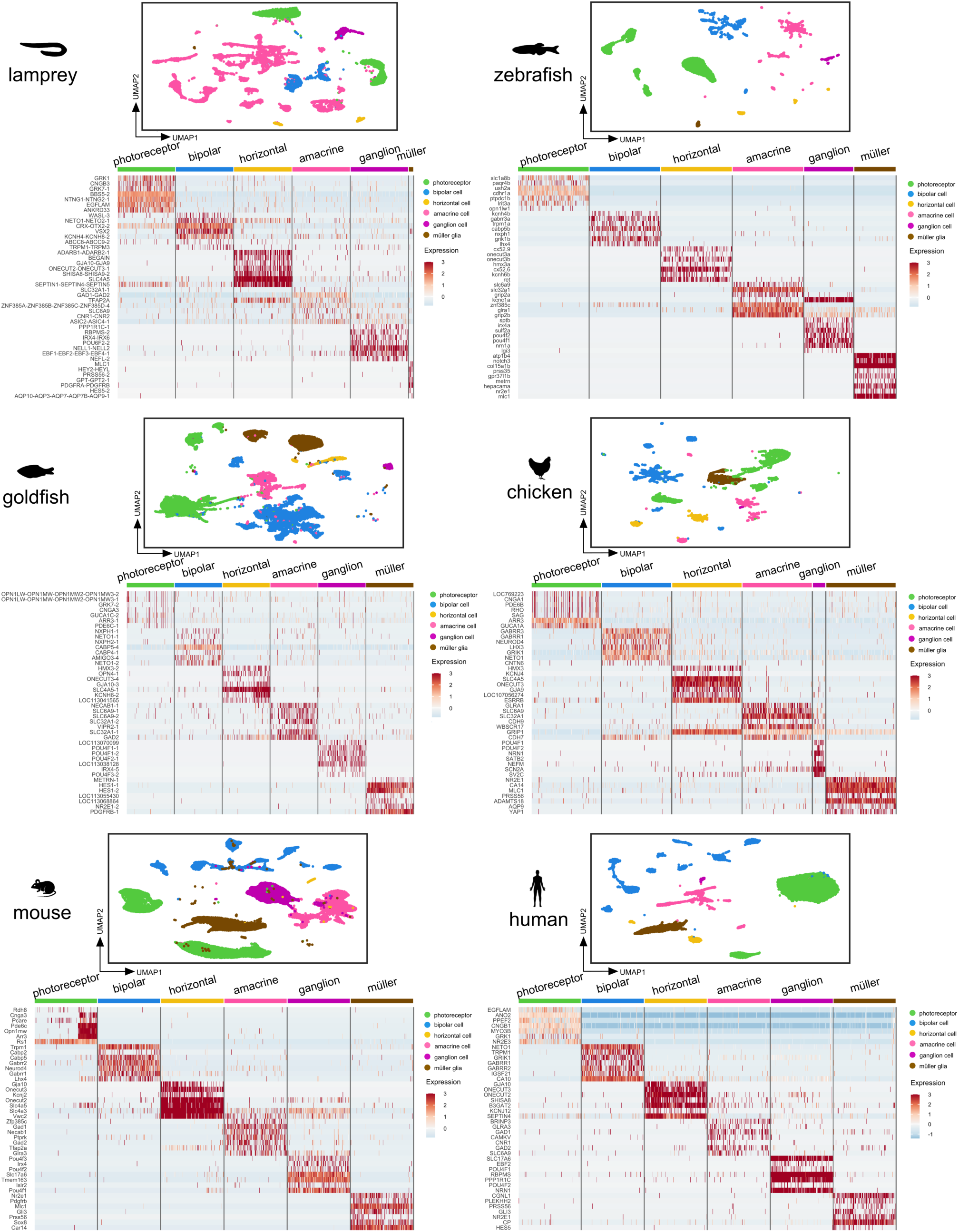
Single-cell expression profiles of retinal cell classes in six species. Retinal cell classes were identified through clustering analysis and visualized using the UMAP method. The heatmap illustrates the expression of selected cell class-enriched marker genes that are conserved across species. Highest-scoring marker genes are shown for each cell class in each species. Gene names in lamprey and goldfish were updated based on the information provided by hierarchical orthogroups (See Methods).

**Supplementary Fig. 2.**
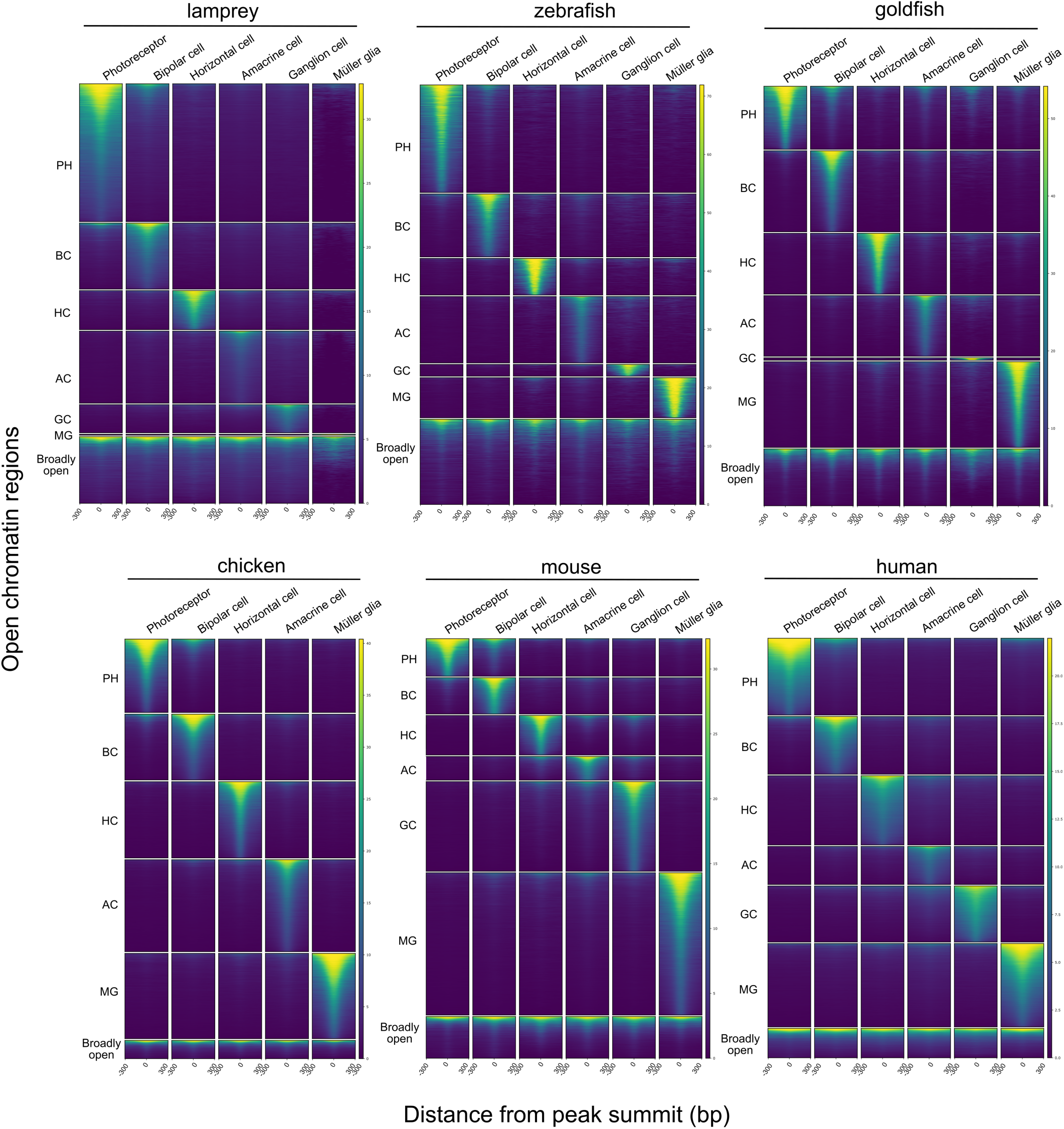
Chromatin accessibility in cell class-enriched and broadly open regions. Columns show pseudo-bulk snATAC-seq profiles for each species, grouped vertically according to cell class or broadly open regions. The color gradient reflects chromatin accessibility from high accessibility (yellow) to low accessibility (dark blue). PH, photoreceptor cells; BC, bipolar cells; HC, horizontal cells; AC, amacrine cells; GC, ganglion cells; MG, Müller glia.

**Supplementary Fig. 3.**
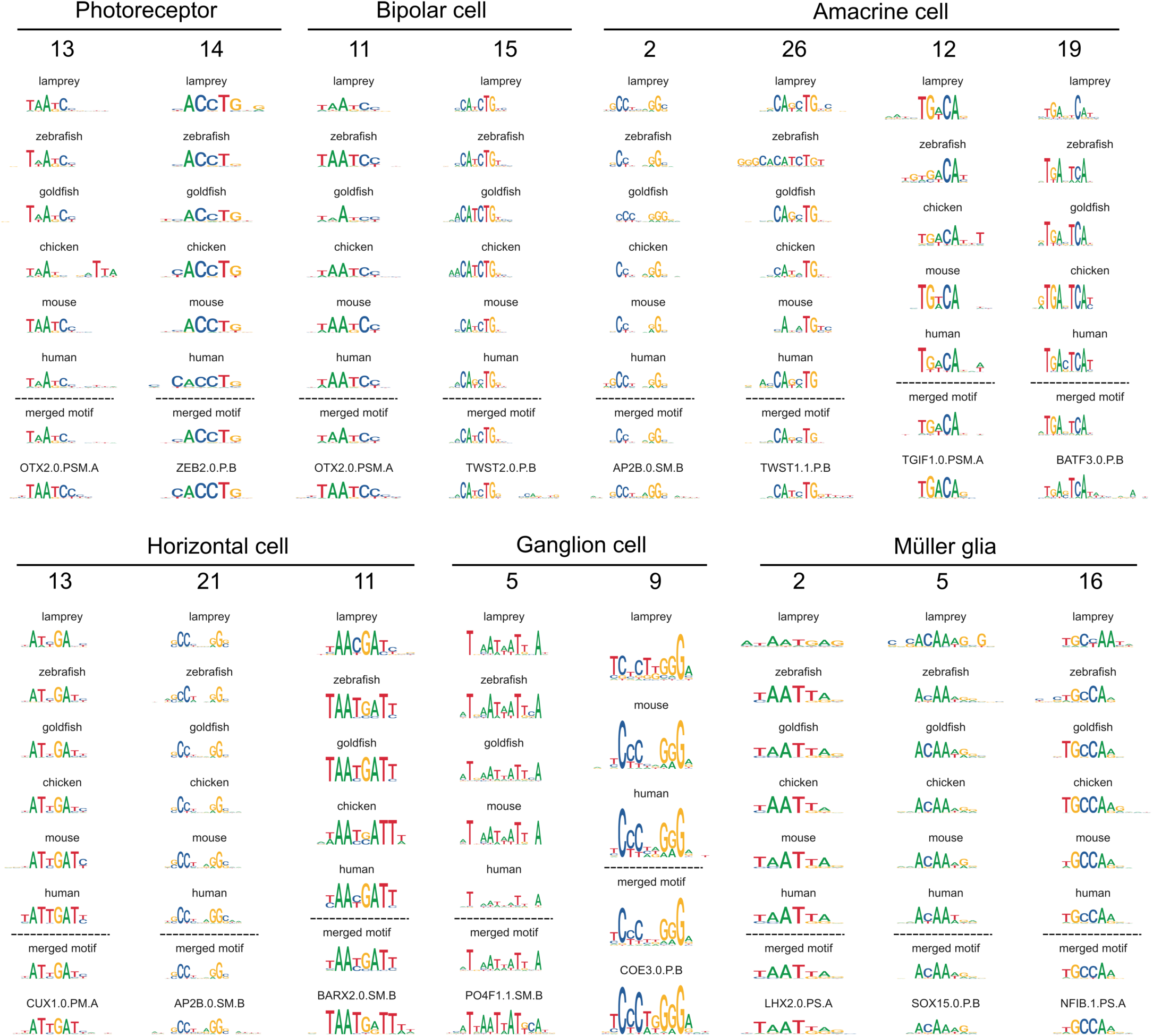
Evolutionarily conserved *cis*-regulatory motifs. The sequence logo for the most significantly enriched motif in each species is shown, along with the logos for a ‘merged’ motif representing the average of the six species motifs and the closest known motif in the *HOCOMOCO* database^23^. The motifs for each cell class are ordered as in Fig. 3d.

**Supplementary Fig. 4.**
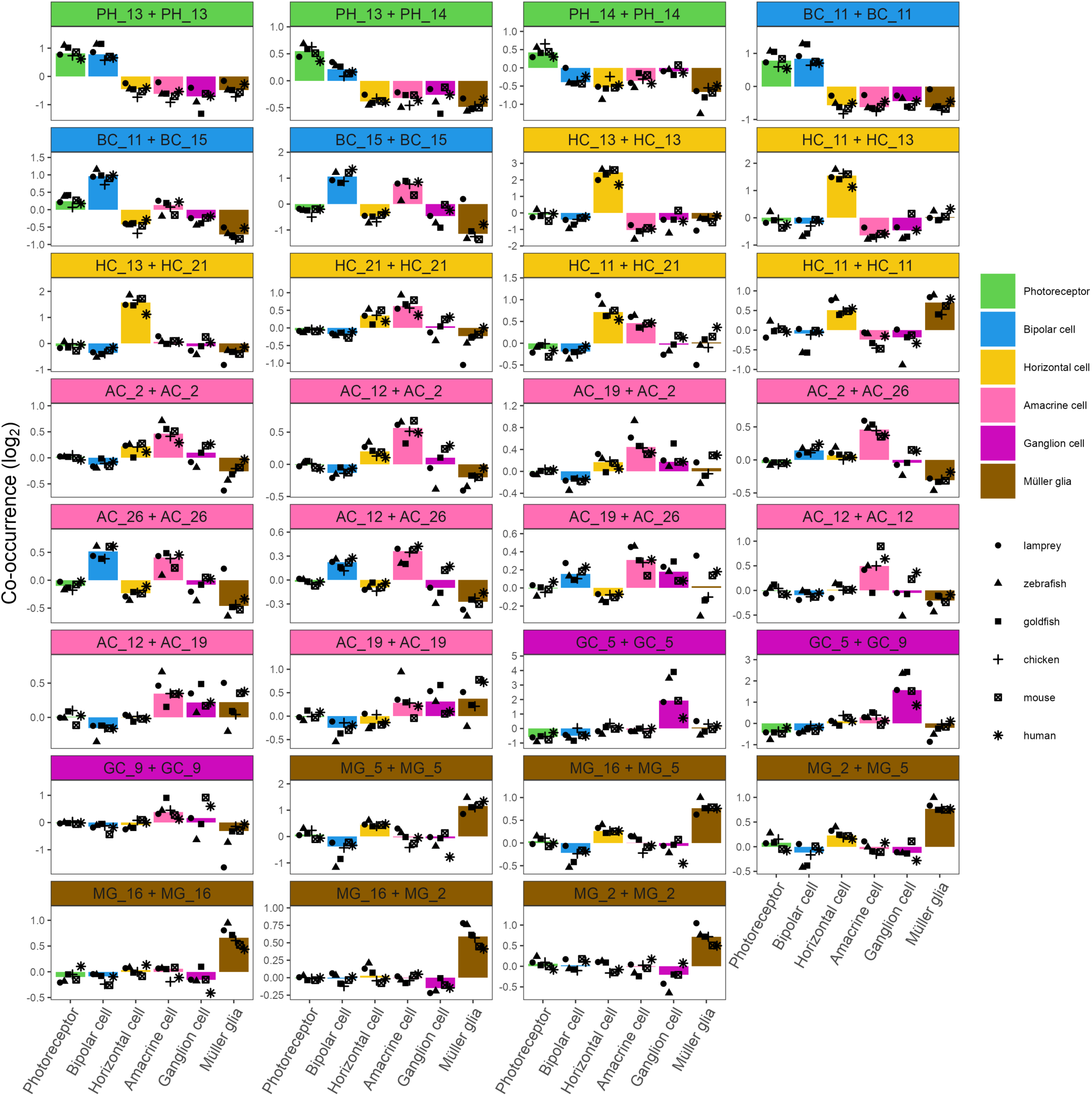
Evolutionarily conserved motif co-occurrence in cell class-enriched open chromatin regions. The graphs illustrate the enrichment of co-occurrence of the indicated motif pairs in cell class-enriched open chromatin regions relative to broadly open regions. The median enrichment across all six species is indicated by the bar, and the values for individual species by symbols.

**Supplementary Fig. 5.**
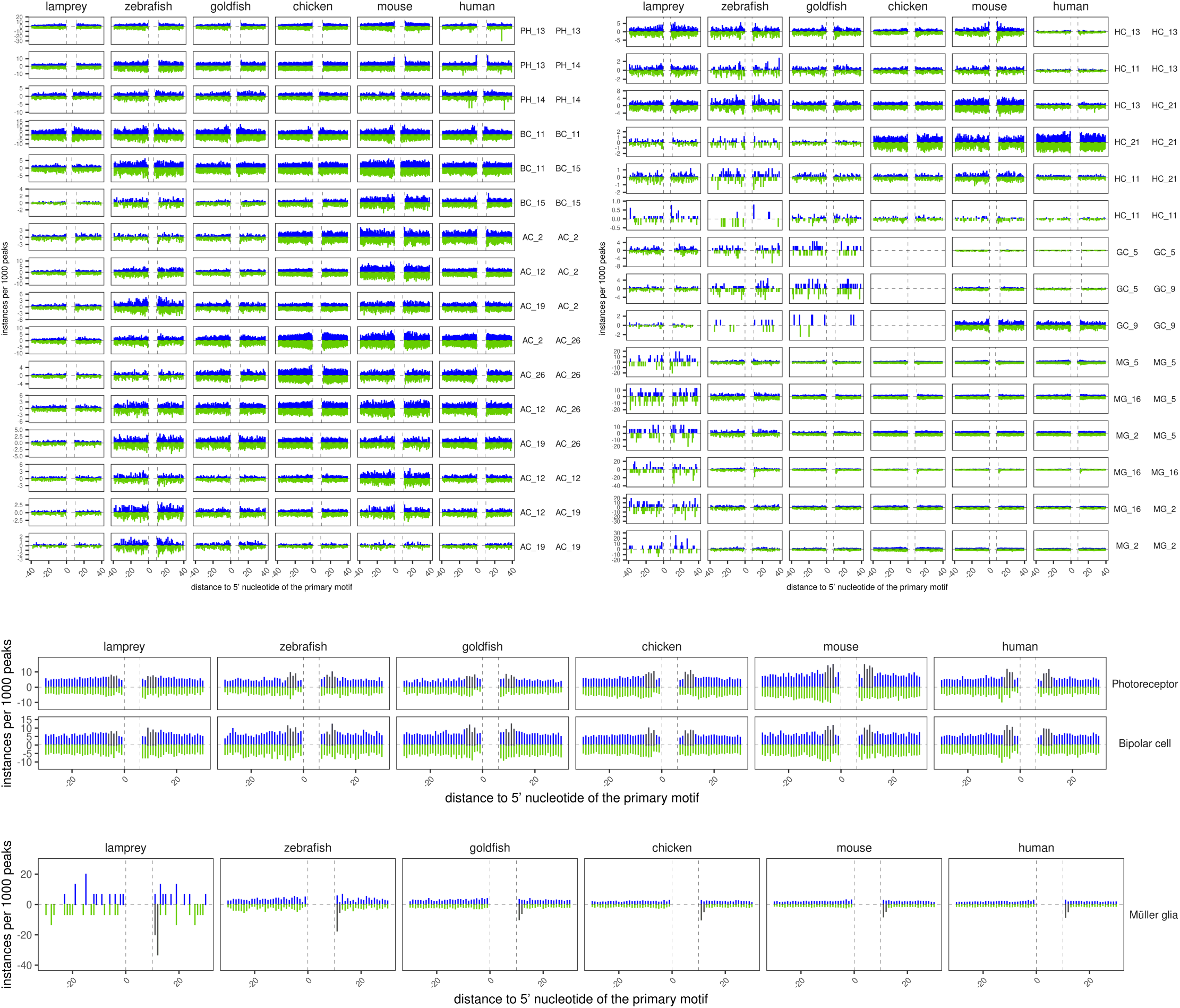
Patterns of spacing and orientation between pairs of evolutionarily conserved motifs. **a,** Each panel shows the strand-specific per nucleotide density of the indicated secondary motif upstream and downstream of the primary motif (located between vertical dotted lines) in the indicated cell class-enriched open chromatin regions. Secondary motifs on the positive strand are shown in blue above the midline. Secondary motifs on the negative strand are shown in green, below the midline. ND, not determined. **b,** Motif spacing patterns between homotypic pairs of monomeric K50 homeodomain-type motifs (i.e., BC_11) in photoreceptor-and bipolar cell-enriched open chromatin regions. Intermotif spacings of 8-11 bp (i.e., approximately one helical turn) are highlighted in gray**. c,** Motif spacing pattern between homotypic pairs of the HMG/SOX-type motif (i.e., MG_16) in Müller glia-enriched open chromatin regions. Intermotif spacings of 4-5 bp, which are characteristic of dimeric SOXE-type binding sites^30^, are highlighted in gray.

**Supplementary Fig. 6.**
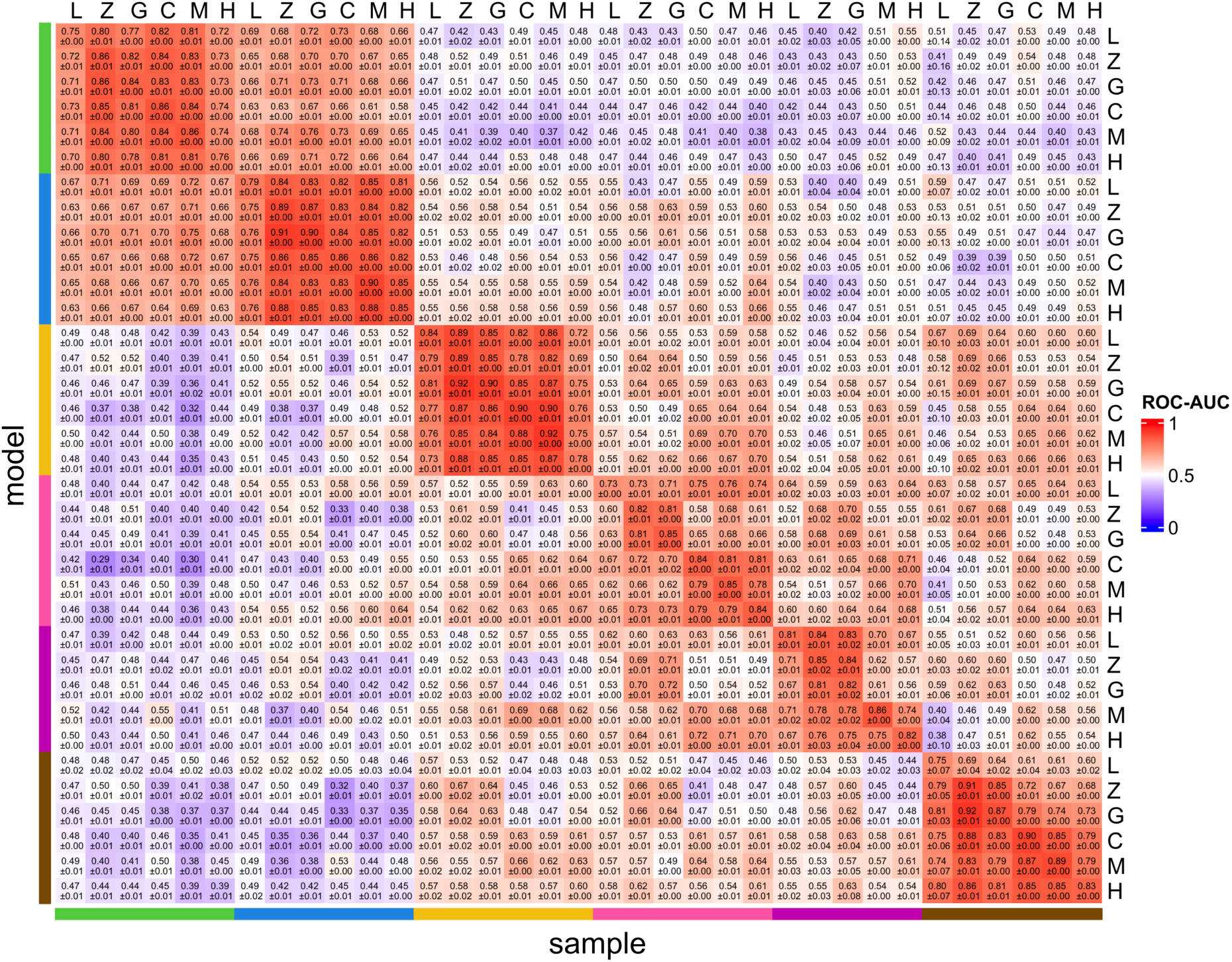
Classificatory performance of gkm-SVM models. The performance of gkm-SVM models in classifying cell class-enriched open chromatin regions is shown. The average and standard deviation of ROC-AUC scores for five gkm-SVM model replicates are indicated within each box. L. lamprey; Z, zebrafish; G, goldfish; C, chicken; M, mouse; H, human.

**Supplementary Fig. 7.**
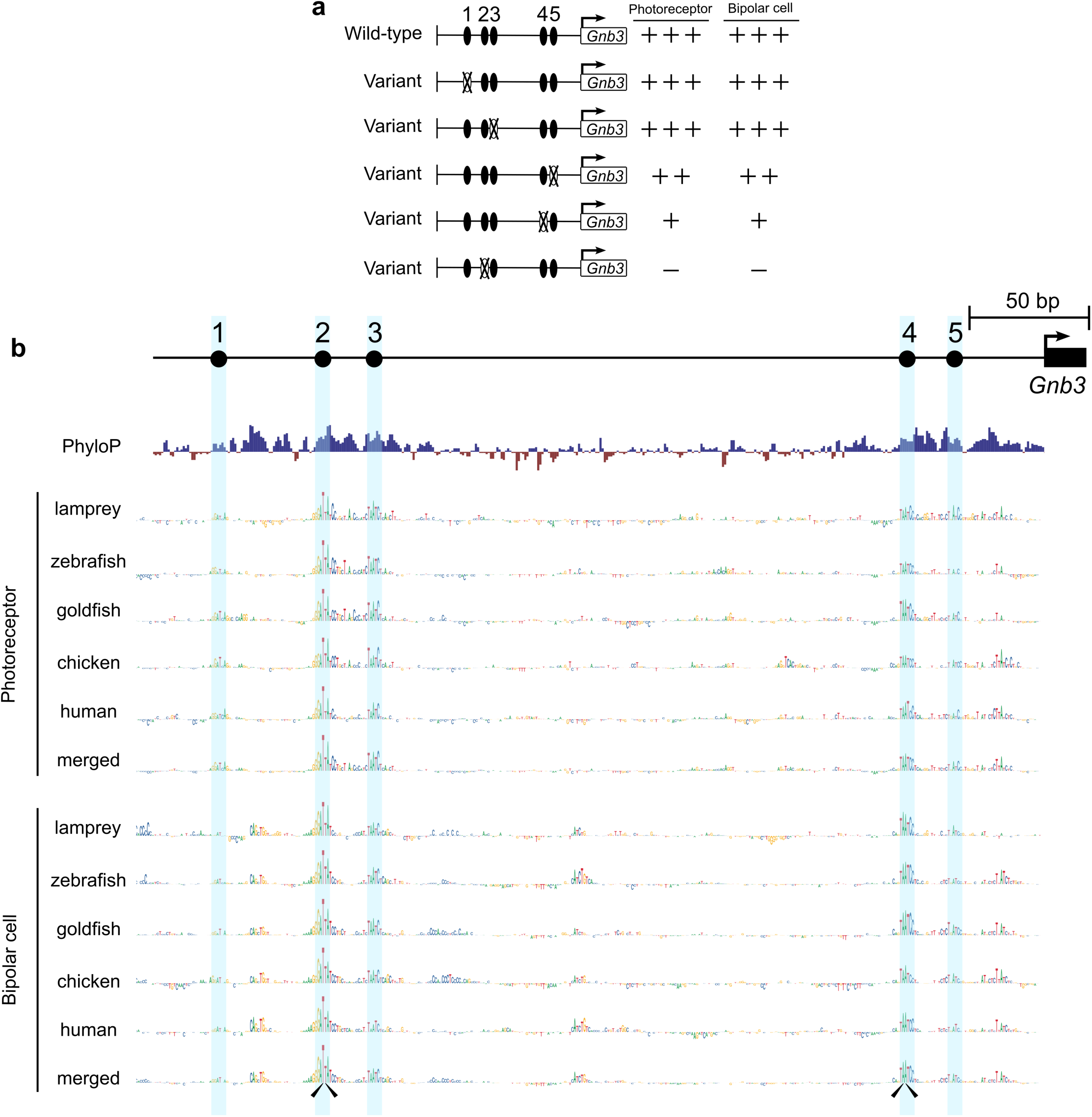
Predicted contribution of individual nucleotides in the mouse *Gnb3* promoter to the overall gkm-SVM model score. **a,** Activity of wild-type and variant mouse *Gnb3* promoter constructs from a previously published electroporation-based reporter assay performed in living mouse retina^11^. The promoter contains five bioinformatically predicted monomeric K50 homeodomain-type binding site motifs numbered 1-5. Inactivating mutations were introduced into each motif in each of the five ‘variant’ constructs. In each case, the core of the motif ‘T<u>AA</u>T’ was changed to ‘T<u>GG</u>T’. +++, strong activity; ++, moderate activity; + weak activity; –, no activity; ×, mutated motif. **b,** The contribution of individual nucleotides to the overall score of the indicated gkm-SVM model is displayed as a sequence logo generated using *GkmExplain*^32^. The height of each nucleotide is proportional to its contribution to the overall score. ‘Merged’ indicates the averaged values across all five models (see Methods). Mutated nucleotides in variant constructs #2 and #4 are indicated with arrowheads.

**Supplementary Fig. 8.**
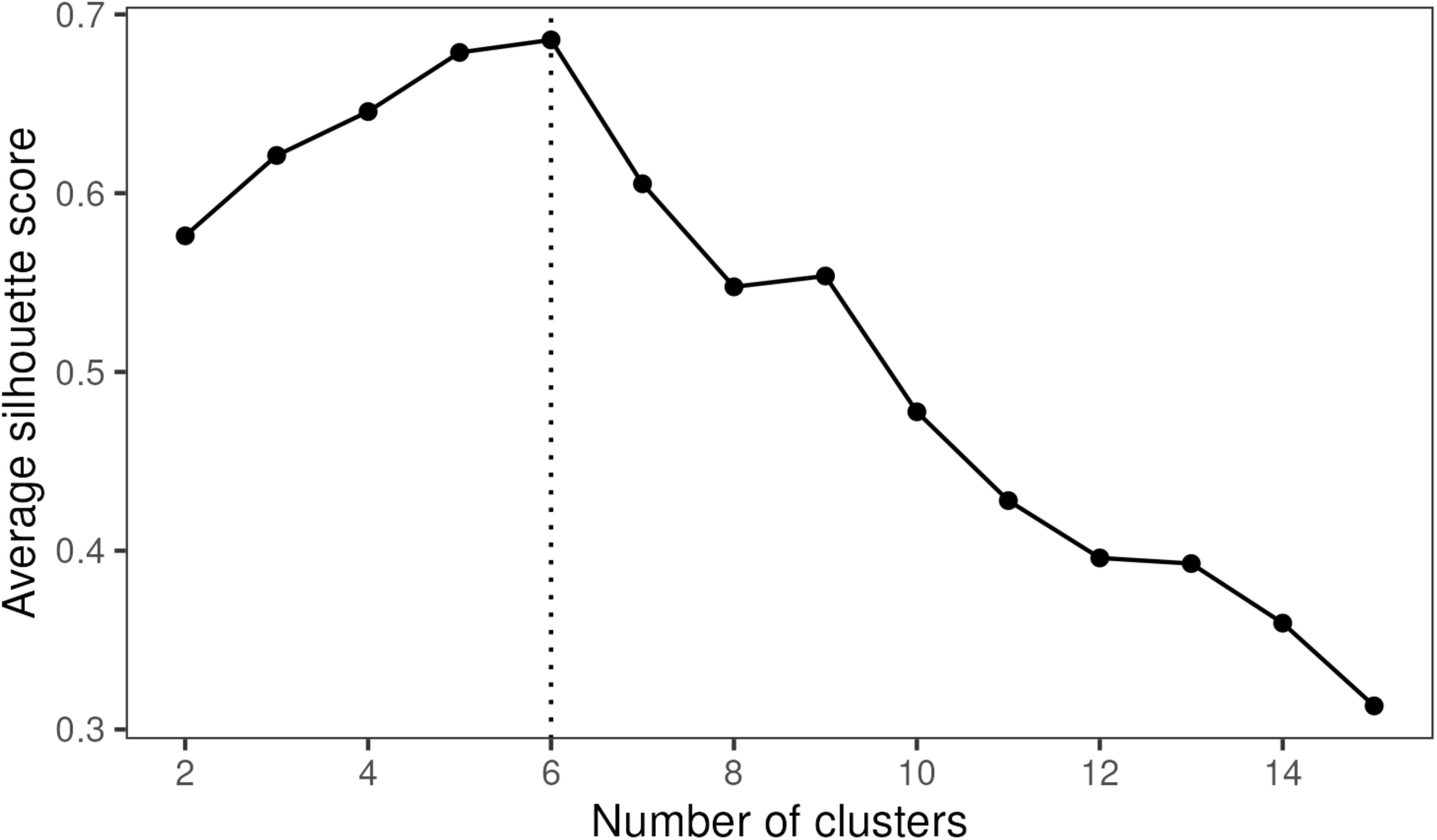
Performance of hierarchical clustering of gkm-SVM models. **Clustering performance** was evaluated using the silhouette score, a measure of cluster cohesion, separation, stability, and robustness. The silhouette scores were averaged across the models within each cluster, and the resulting averages were subsequently averaged across clusters (see Methods).

## Notes

### Competing Interest Statement

The authors have declared no competing interest.

## References

1 Arendt, D., Bertucci, P., Achim, K. & Musser, J. M. Evolution of neuronal types and families. Current opinion in neurobiology 56, 144–152 (2019). 10.1016/j.conb.2019.01.022

2 Arendt, D. et al. The origin and evolution of cell types. Nature Reviews Genetics 17, 744–757 (2016). 10.1038/nrg.2016.127

3 Tosches, M. A. From Cell Types to an Integrated Understanding of Brain Evolution: The Case of the Cerebral Cortex. Annu Rev Cell Dev Biol 37, 495–517 (2021). 10.1146/annurev-cellbio-120319-112654

4 Pomaville, M. B., Sattler, S. M. & Abitua, P. B. A new dawn for the study of cell type evolution. Development (Cambridge, England) 151 (2024). 10.1242/dev.200884

5 Lamb, T. D., Collin, S. P., & Jr, E. N. Evolution of the vertebrate eye: opsins, photoreceptors, retina and eye cup. Nature Reviews Neuroscience 8, 960 (2007). 10.1038/nrn2283

6 Baden, T. The vertebrate retina: a window into the evolution of computation in the brain. Curr Opin Behav Sci 57, 101391 (2024). 10.1016/j.cobeha.2024.101391

7 Hahn, J. et al. Evolution of neuronal cell classes and types in the vertebrate retina. Nature 624, 415–424 (2023). 10.1038/s41586-023-06638-9

8 Fain, G. L. Lamprey vision: Photoreceptors and organization of the retina. Seminars in cell & developmental biology 106, 5–11 (2020). 10.1016/j.semcdb.2019.10.008

9 Wang, J. et al. Molecular Characterization of the Sea Lamprey Retina Illuminates the Evolutionary Origin of Retinal Cell Types. bioRxiv, 2023.2012.2010.571000 (2023). 10.1101/2023.12.10.571000

10 Suzuki, D. G. & Grillner, S. The stepwise development of the lamprey visual system and its evolutionary implications. Biol Rev Camb Philos Soc 93, 1461–1477 (2018). 10.1111/brv.12403

11 Murphy, D. P., Hughes, A. E., Lawrence, K. A., Myers, C. A. & Corbo, J. C. *Cis*-regulatory basis of sister cell type divergence in the vertebrate retina. Elife 8, e48216 (2019). 10.7554/eLife.48216

12 Baden, T. Ancestral photoreceptor diversity as the basis of visual behaviour. Nature Ecology & Evolution, 1–13 (2024). 10.1038/s41559-023-02291-7

13 Baden, T. From water to land: Evolution of photoreceptor circuits for vision in air. PLoS biology 22, e3002422 (2024). 10.1371/journal.pbio.3002422

14 Kim, S. & Wysocka, J. Deciphering the multi-scale, quantitative cis-regulatory code. Molecular cell 83, 373–392 (2023). 10.1016/j.molcel.2022.12.032

15 Levine, M. Transcriptional enhancers in animal development and evolution. Current biology : CB 20, R754–763 (2010). 10.1016/j.cub.2010.06.070

16 Baden, T., Euler, T. & Berens, P. Understanding the retinal basis of vision across species. Nature reviews. Neuroscience 21, 5–20 (2020). 10.1038/s41583-019-0242-1

17 Hughes, A. E., Enright, J. M., Myers, C. A., Shen, S. Q. & Corbo, J. C. Cell type-specific epigenomic analysis reveals a uniquely closed chromatin architecture in mouse rod photoreceptors. Scientific reports 7, 43184 (2017). 10.1038/srep43184

18 Hughes, A. E. O., Myers, C. A. & Corbo, J. C. A massively parallel reporter assay reveals context-dependent activity of homeodomain binding sites in vivo. Genome research 28, 1520–1531 (2018). 10.1101/gr.231886.117

19 Bininda-Emonds, O. R. Fast genes and slow clades: comparative rates of molecular evolution in mammals. Evol Bioinform Online 3, 59–85 (2007).

20 Gibbs, R. A. et al. Genome sequence of the Brown Norway rat yields insights into mammalian evolution. Nature 428, 493–521 (2004). 10.1038/nature02426

21 http://homer.ucsd.edu/homer/motif/index.html.

22 Gupta, S., Stamatoyannopoulos, J. A., Bailey, T. L. & Noble, W. S. Quantifying similarity between motifs. Genome biology 8, R24 (2007). 10.1186/gb-2007-8-2-r24

23 Kulakovskiy, I. V. et al. HOCOMOCO: towards a complete collection of transcription factor binding models for human and mouse via large-scale ChIP-Seq analysis. Nucleic acids research 46, D252–d259 (2018). 10.1093/nar/gkx1106

24 Freund, C. L. et al. Cone-rod dystrophy due to mutations in a novel photoreceptor-specific homeobox gene (CRX) essential for maintenance of the photoreceptor. Cell 91, 543–553 (1997).

25 Furukawa, T., Morrow, E. M. & Cepko, C. L. *Crx*, a novel *otx*-like homeobox gene, shows photoreceptor-specific expression and regulates photoreceptor differentiation. Cell 91, 531–541 (1997).

26 Bassett, E. A. & Wallace, V. A. Cell fate determination in the vertebrate retina. Trends in neurosciences 35, 565–573 (2012). 10.1016/j.tins.2012.05.004

27 Petridou, E. & Godinho, L. Cellular and Molecular Determinants of Retinal Cell Fate. Annu. Rev. Vis. Sci. 8, 79–99 (2022). 10.1146/annurev-vision-100820-103154

28 Emms, D. M. & Kelly, S. OrthoFinder: phylogenetic orthology inference for comparative genomics. Genome biology 20, 238 (2019). 10.1186/s13059-019-1832-y

29 Corbo, J. C. et al. CRX ChIP-seq reveals the cis-regulatory architecture of mouse photoreceptors. Genome research 20, 1512–1525 (2010). 10.1101/gr.109405.110

30 Huang, Y.-H., Jankowski, A., Cheah, K. S. E., Prabhakar, S. & Jauch, R. SOXE transcription factors form selective dimers on non-compact DNA motifs through multifaceted interactions between dimerization and high-mobility group domains. Scientific reports 5, 10398 (2015). 10.1038/srep10398

31 Ghandi, M., Lee, D., Mohammad-Noori, M. & Beer, M. A. Enhanced Regulatory Sequence Prediction Using Gapped k-mer Features. PLoS computational biology 10, e1003711 (2014). 10.1371/journal.pcbi.1003711

32 Shrikumar, A., Prakash, E. & Kundaje, A. GkmExplain: fast and accurate interpretation of nonlinear gapped k-mer SVMs. Bioinformatics (Oxford, England) 35, i173–i182 (2019). 10.1093/bioinformatics/btz322

33 Wunderlich, Z. & Mirny, L. A. Different gene regulation strategies revealed by analysis of binding motifs. Trends in Genetics 25, 434–440 (2009). 10.1016/j.tig.2009.08.003

34 Hobert, O. Regulatory logic of neuronal diversity: terminal selector genes and selector motifs. Proc Natl Acad Sci U S A 105, 20067–20071 (2008). 10.1073/pnas.0806070105

35 Mears, A. J. et al. Nrl is required for rod photoreceptor development. Nature genetics 29, 447–452 (2001).

36 Kram, Y. A., Mantey, S. & Corbo, J. C. Avian cone photoreceptors tile the retina as five independent, self-organizing mosaics. PloS one 5, e8992 (2010). 10.1371/journal.pone.0008992

37 Coolen, M. et al. Phylogenomic analysis and expression patterns of large Maf genes in Xenopus tropicalis provide new insights into the functional evolution of the gene family in osteichthyans. Development genes and evolution 215, 327–339 (2005). 10.1007/s00427-005-0476-y

38 Ochi, H. et al. Temporal expression of L-Maf and RaxL in developing chicken retina are arranged into mosaic pattern. Gene expression patterns : GEP 4, 489–494 (2004). 10.1016/j.modgep.2004.03.005

39 Enright, J. M., Lawrence, K. A., Hadzic, T. & Corbo, J. C. Transcriptome profiling of developing photoreceptor subtypes reveals candidate genes involved in avian photoreceptor diversification. The Journal of comparative neurology (2014). 10.1002/cne.23702

40 Kim, J.-W. et al. Recruitment of Rod Photoreceptors from Short-Wavelength-Sensitive Cones during the Evolution of Nocturnal Vision in Mammals. Dev Cell 37, 520–532 (2016). 10.1016/j.devcel.2016.05.023

41 Oel, A. P. et al. Nrl is dispensable for specification of rod photoreceptors in adult zebrafish despite its deeply conserved requirement earlier in ontogeny. Iscience 23, 101805 (2020). 10.1016/j.isci.2020.101805

42 Liu, F. et al. Rod genesis driven by mafba in an nrl knockout zebrafish model with altered photoreceptor composition and progressive retinal degeneration. PLoS genetics 18, e1009841 (2022). 10.1371/journal.pgen.1009841

43 Kerppola, T. K. & Curran, T. A conserved region adjacent to the basic domain is required for recognition of an extended DNA binding site by Maf/Nrl family proteins. Oncogene 9, 3149–3158 (1994).

44 Guide for the Care and Use of Laboratory Animals. 8th edn, (The National Academies Press, 2011).

45 Li, H. Minimap2: pairwise alignment for nucleotide sequences. Bioinformatics (Oxford, England) 34, 3094–3100 (2018). 10.1093/bioinformatics/bty191

46 Kon, T. et al. Single-cell transcriptomics of the goldfish retina reveals genetic divergence in the asymmetrically evolved subgenomes after allotetraploidization. Commun Biology 5, 1404 (2022). 10.1038/s42003-022-04351-3

47 Lyu, P. et al. Gene regulatory networks controlling temporal patterning, neurogenesis, and cell-fate specification in mammalian retina. Cell reports 37, 109994 (2021). 10.1016/j.celrep.2021.109994

48 Lyu, P. et al. Common and divergent gene regulatory networks control injury-induced and developmental neurogenesis in zebrafish retina. Nat Commun 14, 8477 (2023). 10.1038/s41467-023-44142-w

49 Wang, S. K. et al. Single-cell multiome of the human retina and deep learning nominate causal variants in complex eye diseases. Cell Genom 2 (2022). 10.1016/j.xgen.2022.100164

50 Granja, J. M. et al. ArchR is a scalable software package for integrative single-cell chromatin accessibility analysis. Nature genetics 53, 403–411 (2021). 10.1038/s41588-021-00790-6

51 Hao, Y. et al. Integrated analysis of multimodal single-cell data. Cell 184, 3573–3587.e3529 (2021). 10.1016/j.cell.2021.04.048

52 Stuart, T. et al. Comprehensive integration of single cell data. bioRxiv, 460147 (2018). 10.1101/460147

53 Stuart, T., Srivastava, A., Madad, S., Lareau, C. A. & Satija, R. Single-cell chromatin state analysis with Signac. Nature methods 18, 1333–1341 (2021). 10.1038/s41592-021-01282-5

54 Yamagata, M., Yan, W. & Sanes, J. R. A cell atlas of the chick retina based on single-cell transcriptomics. Elife 10 (2021). 10.7554/eLife.63907

55 Clark, B. S. et al. Single-Cell RNA-Seq Analysis of Retinal Development Identifies NFI Factors as Regulating Mitotic Exit and Late-Born Cell Specification. Neuron 102, 1111–1126.e1115 (2019). 10.1016/j.neuron.2019.04.010

56 Korsunsky, I. et al. Fast, sensitive and accurate integration of single-cell data with Harmony. Nature methods 16, 1289–1296 (2019). 10.1038/s41592-019-0619-0

57 Li, J. et al. Comprehensive single-cell atlas of the mouse retina. Iscience 27, 109916 (2024). 10.1016/j.isci.2024.109916

58 Yanai, I. et al. Genome-wide midrange transcription profiles reveal expression level relationships in human tissue specification. *Bioinformatics (Oxford*, England) 21, 650–659 (2005). 10.1093/bioinformatics/bti042

59 Ramírez, F. et al. deepTools2: a next generation web server for deep-sequencing data analysis. Nucleic acids research 44, W160–165 (2016). 10.1093/nar/gkw257

60 Nassar, L. R. et al. The UCSC Genome Browser database: 2023 update. Nucleic acids research 51, D1188–d1195 (2023). 10.1093/nar/gkac1072

61 Pracana, R., Priyam, A., Levantis, I., Nichols, R. A. & Wurm, Y. The fire ant social chromosome supergene variant Sb shows low diversity but high divergence from SB. Mol Ecol 26, 2864–2879 (2017). 10.1111/mec.14054

62 Hedges, S. B., Marin, J., Suleski, M., Paymer, M. & Kumar, S. Tree of life reveals clock-like speciation and diversification. Molecular biology and evolution 32, 835–845 (2015). 10.1093/molbev/msv037

63 Vierstra, J. et al. Global reference mapping of human transcription factor footprints. Nature 583, 729–736 (2020). 10.1038/s41586-020-2528-x

64 http://meme-suite.org/.

65 Ou, J., Wolfe, S. A., Brodsky, M. H. & Zhu, L. J. motifStack for the analysis of transcription factor binding site evolution. Nature methods 15, 8–9 (2018). 10.1038/nmeth.4555

66 Rosanova, A., Colliva, A., Osella, M. & Caselle, M. Modelling the evolution of transcription factor binding preferences in complex eukaryotes. Scientific reports 7, 7596 (2017). 10.1038/s41598-017-07761-0

67 Lee, D. LS-GKM: a new gkm-SVM for large-scale datasets. *Bioinformatics (Oxford*, England) 32, 2196–2198 (2016). 10.1093/bioinformatics/btw142

68 Sing, T., Sander, O., Beerenwinkel, N. & Lengauer, T. ROCR: visualizing classifier performance in R. *Bioinformatics (Oxford*, England) 21, 3940–3941 (2005). 10.1093/bioinformatics/bti623

